# Outer membrane vesicles as realistic models of bacterial membranes in interaction studies by Surface Plasmon Resonance

**DOI:** 10.1101/2023.07.07.548064

**Authors:** Maxim S. Bril’kov, Victoria Stenbakk, Martin Jakubec, Terje Vasskog, Tone Kristoffersen, Jorunn Pauline Cavanagh, Johanna U. Ericson, Johan Isaksson, Gøril Eide Flaten

## Abstract

One way to mitigate the ongoing antimicrobial resistance crisis is to discover and develop new classes of antibiotics. As all antibiotics at some point needs to either cross or interact with the bacterial membrane, there is a need for representative models of bacterial membranes and efficient methods to characterize the interactions to novel antimicrobials – both to generate new knowledge and to screen compound libraries. Since the bacterial cell envelope is a complex assembly of lipids, lipopolysaccharides, membrane proteins and other components, constructing realistic synthetic liposome-based models of the membrane is both difficult and expensive.

We here propose to let the bacteria do the hard work for us. Outer membrane vesicles (OMVs) are naturally secreted by Gram-negative bacteria, playing a role in communication between bacteria, as virulence factors, molecular transport or being a part of the antimicrobial resistance mechanism. OMVs consist of the bacterial outer membrane and thus inherit many components and properties of the native outer cell envelope. In this work we have isolated and characterized OMVs from *E. coli* mutant strains and clinical isolates of the ESKAPE members *Klebsiella pneumoniae, Acinetobacter baumannii* and *Pseudomonas aeruginosa*. The OMVs were shown to be representative models for the bacterial membrane in terms of lipid composition with strain specific variations. The OMVs were further used to probe the interactions between OMV and antimicrobial peptides (AMPs) as model compounds by Surface Plasmon Resonance (SPR) and provide proof-of-principle that OMVs can be used as an easily accessible and highly realistic model for the bacterial surface in interaction studies. This further enables direct monitoring of the effect of induction by antibiotics, or the response to host-pathogen interactions.

## Introduction

The development of antimicrobial resistance (AMR) is a serious threat to public health and the main obstacle to successful treatment of infectious diseases. AMR caused 4.95 million deaths already in 2019 and the numbers are growing (1, 2). The antimicrobial resistant bacteria responsible for more than 15 % of hospital acquired infections are classified as global high priority pathogens by the World Health Organization (WHO). The Gram-negative bacteria *K. pneumoniae, A. baumannii* and *P. aeruginosa* are on top of the list and are thus representing the strains of critical priority (3, 4).

To fight the ongoing antimicrobial resistance crisis effort is put into the discovery and optimization of new antimicrobial hit compounds. As most aspiring antibiotics at some point need to either cross or interact with the bacterial membrane, there is a need for cost-effective native like models of bacterial membranes and efficient methods to characterize the interactions with novel antimicrobials (5).

The cell envelope of Gram-negative bacteria is a complex structure consisting of inner and outer membranes. The inner membrane is mainly composed of phospholipids, while the outer membrane is asymmetric and composed mainly of phospholipids (PLs) at the inner leaflet and lipopolysaccharides (LPS) at the outer leaflet. The major structural unit of LPS is lipid A, and the net charge of LPS is negative. During infection, or in response to antibiotic treatment, bacteria can modify the lipid A structure to alter the surface charge which could lead to reduced binding of cationic antimicrobials such as colistin (6-10). The periplasmic space between the inner and outer membranes is enriched in peptidoglycans. The cell envelope is also the site for various proteins such as ion pumps and channels, receptors and membrane bound enzymes (11-13). Simple synthetic lipids in the form of liposomes, supported lipid bilayers, or nanodiscs are often utilized to investigate new antimicrobial compounds. However, they lack the intricate complexity observed in the native cell envelope (14-20). Incorporation of all native outer membrane components into synthetic model liposomes is possible but a laborious and expensive process (21-25). There is thus a need for affordable but realistic model systems that can accurately mimic the bacterial membrane.

Outer membrane vesicles (OMVs) are formed mainly from outer membranes and maintain similar composition containing PLs, LPS and OM-proteins, and thus, can represent naturally assembled nano-models of the outer membranes (12, 26). For bacteria, the OMVs play an extensive role in physiological and pathogenic processes in for example communication, biofilm formation, antibiotic resistance, and molecular transport. As a part of the bacterial defense mechanism, OMVs serve as a decoy, absorbing and neutralizing membrane active agents, including bacteriophages. OMVs can also be a central part of a disposal mechanism when a bacteria can get rid of the affected membranes (27, 28). These features of OMVs on their own make them competent object for drug research. OMVs are ideal candidates to serve as the outer membrane model for the bacterial strain it is produced from.

The aim of this work was to explore the applicability of native OMVs as bacterial membrane models in interaction studies. To achieve this aim, OMVs secreted by overproducing *E. coli* Δ*tolA* mutant (29), LPS deficient *E. coli* NR698 strain (30, 31) and clinical isolates of multi-resistant strains of critical importance *A. baumannii* K47-42, *P. aeruginosa* K34-7 and *K. pneumoniae* K47-25 (32) were isolated and characterized. The strains of clinical isolates were collected at University Hospital of Northern Norway and have shown to be resistant to beta-lactam class antibiotics by production of beta-lactamase (32). *K. pneumoniae* and *A. baumannii* strains are also colistin resistant, being able to modify lipid A in the outer membrane (32). The kinetic rates of interaction between the OMVs and membrane active compounds were measured by surface plasmon resonance (SPR). A test panel of cyclic antimicrobial peptides (AMPs). AMPs as model system is due to them receiving increasing attention in the fight against AMR, as well as their different mode of actions leading to destabilization of bacterial membranes and bacteria death (33-36). A test panel of five cyclic antimicrobial peptides (AMP)s with various distributions of charged and hydrophobic amino acids, known to interact with bacterial membranes was used (37-42). This set of peptides serves as a model itself, allowing to study how the charge distribution affects the interactions and has previously been used by our lab to explore kinetics of binding towards synthetic liposomes, nanodiscs and to series of *E. coli* strains (40, 42).

## Results and Discussion

In this work we aimed to apply OMVs as a realistic model of the native bacterial membrane for interaction studies by SPR. Antimicrobial peptides were used as test compounds to establish proof of principle. An *E. coli* strain with a deletion of the *tolA* gene, which results in higher OMV yields (29), was used as a starting point to establish the production- and isolation protocols. The LPS deficient *E. coli* strain NR698 (30, 31) as well as clinical isolates of multidrug resistant strains of *A. baumannii, P. aeruginosa* and *K. pneumoniae* were further selected. The latter belong to a group of strains of critical priority listed by WHO (4). At the first stage, the OMVs were characterized in terms of morphology and lipid composition and compared to bacterial lipid isolates. The OMVs were then utilized directly for interaction studies with the selected test compounds by SPR. In addition to the OMVs, liposomes were prepared from lipids isolated from *E. coli* Δ*tolA* and used as control vesicles in this study.

### Characterization of native OMVs

#### Morphology and zeta potential of isolated OMVs

All strains, except the LPS deficient *E. coli* NR698, produced OMVs in sufficient amounts and quality. Isolated OMVs were visualized by transmission electron microscopy (TEM) to assess morphology and quality of the sample. Figure 1 shows micrographs of representative samples of OMVs from all strains, except *E. coli* NR698. The samples appear to be clear of contamination with only minor traces of cell debris, confirming the successful isolation of the OMVs. The vesicles of all samples were observed to be spherical with a visible double layer. The micrograph of the OMVs from *P. aeruginosa* displayed aggregated vesicles (Figure 1D), a feature commonly observed for this strain in this work. For the LPS deficient *E. coli* NR698 strain, although some OMVs were present, the sample was dominated by deformed OMVs (Figure S1). The strain has mutations causing deficiency of LPS in the outer membrane, resulting in possibly higher content of phospholipids (30, 31). This sample represents the limits of OMVs application and experimental design regarding genetic alterations affecting the properties of the outer membranes. Due to the inadequate quality of the isolated OMVs, this strain was not further pursued.

**Figure 1.**
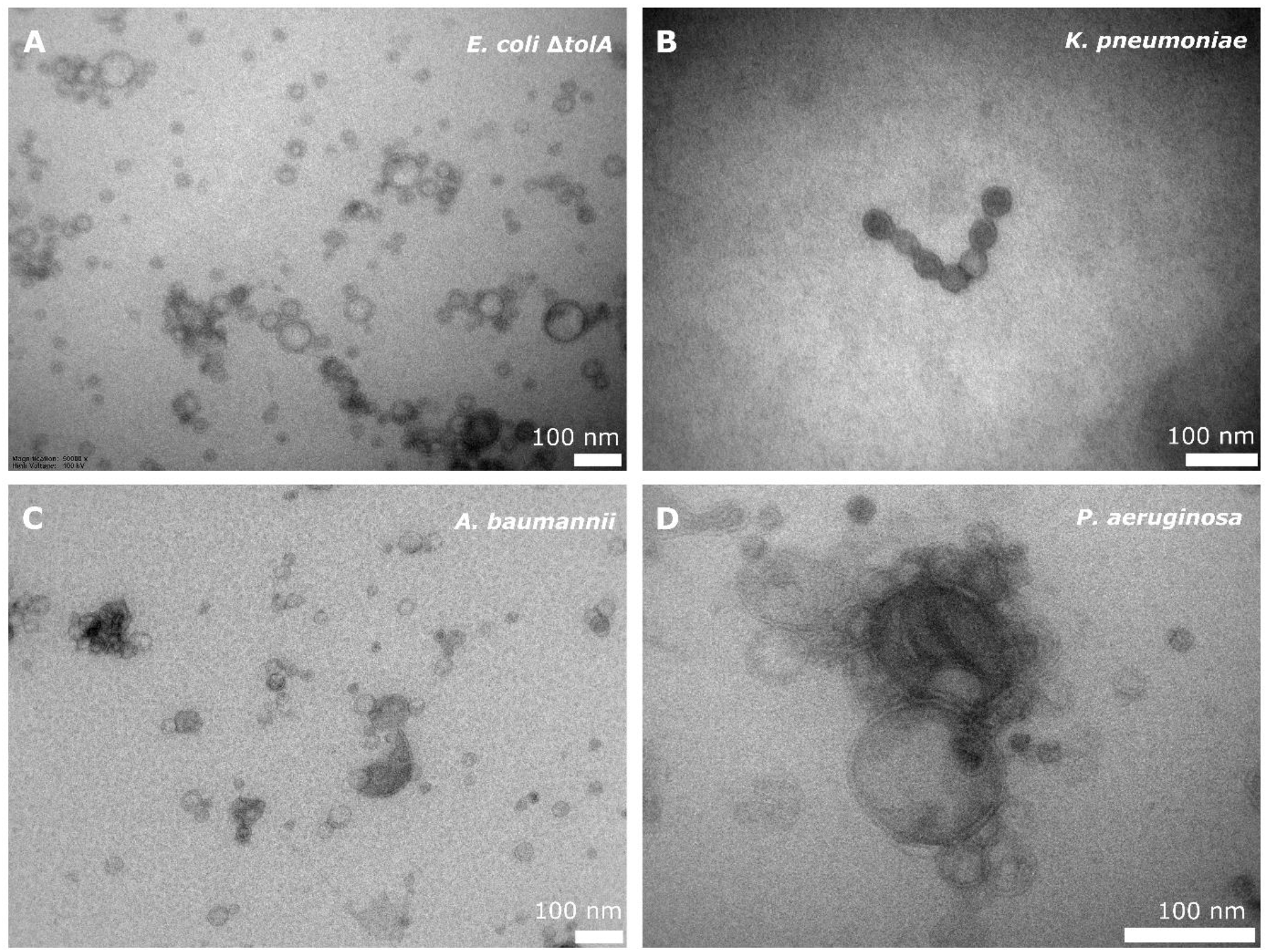
TEM micrograph of OMVs secreted by *E. coli* Δ*tolA* (**A**), *K. pneumoniae* (**B**), *A. baumannii* (**C**) (50,000x magnification), and *P. aeruginosa* (**D**) (80,000x magnification). 100 nm scalebar is presented at the bottom right corner of each micrograph.

The average diameters were determined to be in the range from 194.8 ± 1.0 nm for *K. pneumoniae*, 175.0 ± 1.3 nm for *A. baumannii* and 143.3 ± 1.1 nm for *P. aeruginosa*, to 82.7 ± 1.3 nm for *E. coli* Δ*tolA* (Figure 2A). When comparing the zeta potentials of OMVs and bacterial cells (Figure2B and Table S1), the values are found to be significantly lower for OMVs in all the cases except *P. aeruginosa, w*here no significant difference was observed (T-test, p > 0.05). The lowest zeta potential of −14.4 ± 0.6 mV was measured for OMVs from *A. baumannii*. OMVs from *K. pneumoniae* and *E. coli* Δ*tolA* showed similar values for zeta potentials of −9.9 ± 0.3 mV and −9.4 ± 0.8 mV respectively. The zeta potential of the isolated OMVs from *P. aeruginosa* was measured to be −3.90 ± 0.6 mV, which suggests a less stable dispersion due to less repulsion between the vesicles, and could, thus, explain the aggregation observed (Figure 1D). Liposomes prepared from *E. coli* Δ*tolA* lipid isolate were used as a control in our studies and were also characterized according to size distribution and zeta potential. They showed an average size of 139.9 ± 1.6 nm and had a significantly lower zeta potential, −17.8 ± 0.5 mV, compared to both OMVs produced by the same bacteria, and the whole bacteria itself. This is an indication that the OMVs are a better representation of the bacterial surface than liposomes prepared from lipid isolates. The size of the OMVs isolated from *E. coli* Δ*tolA* was reanalyzed after 1 month of storage at 4 °C. No significant change in size was observed, suggesting that the OMVs were stable for at least 1 month. All samples were analyzed in triplicates.

**Figure 2.**
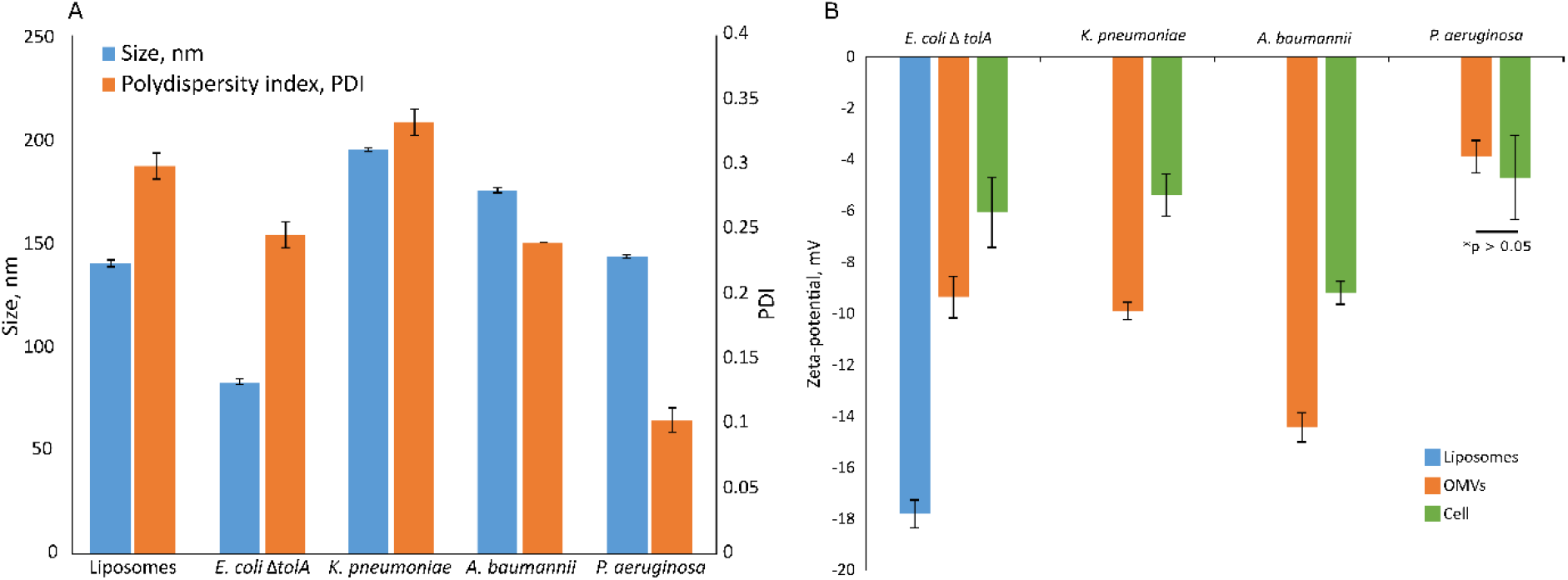
Summary of size and polydispersity index (PDI) (A) and zeta-potential measurements (B) for isolated OMVs in comparison to zeta-potential of the bacterial cells. The sample of liposomes prepared from *E. coli* Δ*tolA* lipid isolate is also included. Samples were held in 10 mM TRIS buffer (100 mM NaCl and 5 mM KCl, pH 8.0) at 25 °C. All samples were analyzed in triplicates.

#### Lipid composition of native bacterial membranes

The compositions of the bacteria lipid isolates were quantified by ^31^P NMR, and a qualitative fingerprint was acquired by MS. The corresponding fingerprint was subsequently acquired for the corresponding secreted OMVs from the same strains.

The results from the ^31^P NMR studies on the whole bacteria lipid isolate are shown in Figure 3A. Phosphatidylethanolamine (PE) was determined to be the dominant phospholipid of the lipid isolates, ranging from 50 – 52 % (for *A. baumanii* and *K. pneumoniae*) up to 62 – 64 % (for *P. aeruginosa* and *E. coli* Δ*tolA*) of the total phospholipid content. The cardiolipin (CL) content was found to be 7 % for *K. pneumoniae*, 10 – 11 % for *P. aeruginosa* and *E. coli* Δ*tolA* and up to 21 % for *A. baumannii*. Lyso-phosphatidylethanolamine (LPE) was determined to be at 24 % and 17 % levels for *K. pneumoniae* and *A. baumannii* respectively, while no LPE was detected in *P. aeruginosa* and *E. coli* Δ*tolA*. Phosphatidylcholine (PC) contributed to 3, 9 and 29 % in *A. baumannii, P. aeruginosa* and *E. coli* Δ*tolA* respectively, while it was not detected in *K. pneumoniae*. Phosphatidylglycerol (PG) contributed to 9 % of the total phospholipids in *E. coli* Δ*tolA* and *A. baumannii*, and up to 17 % for *K. pneumoniae* and *P. aeruginosa. In* conclusion, the PE lipid was the major lipid of the studied bacterial membranes, together with a handful of other phospholipids in smaller quantities, in agreement with previous reports (43-46).

**Figure 3.**
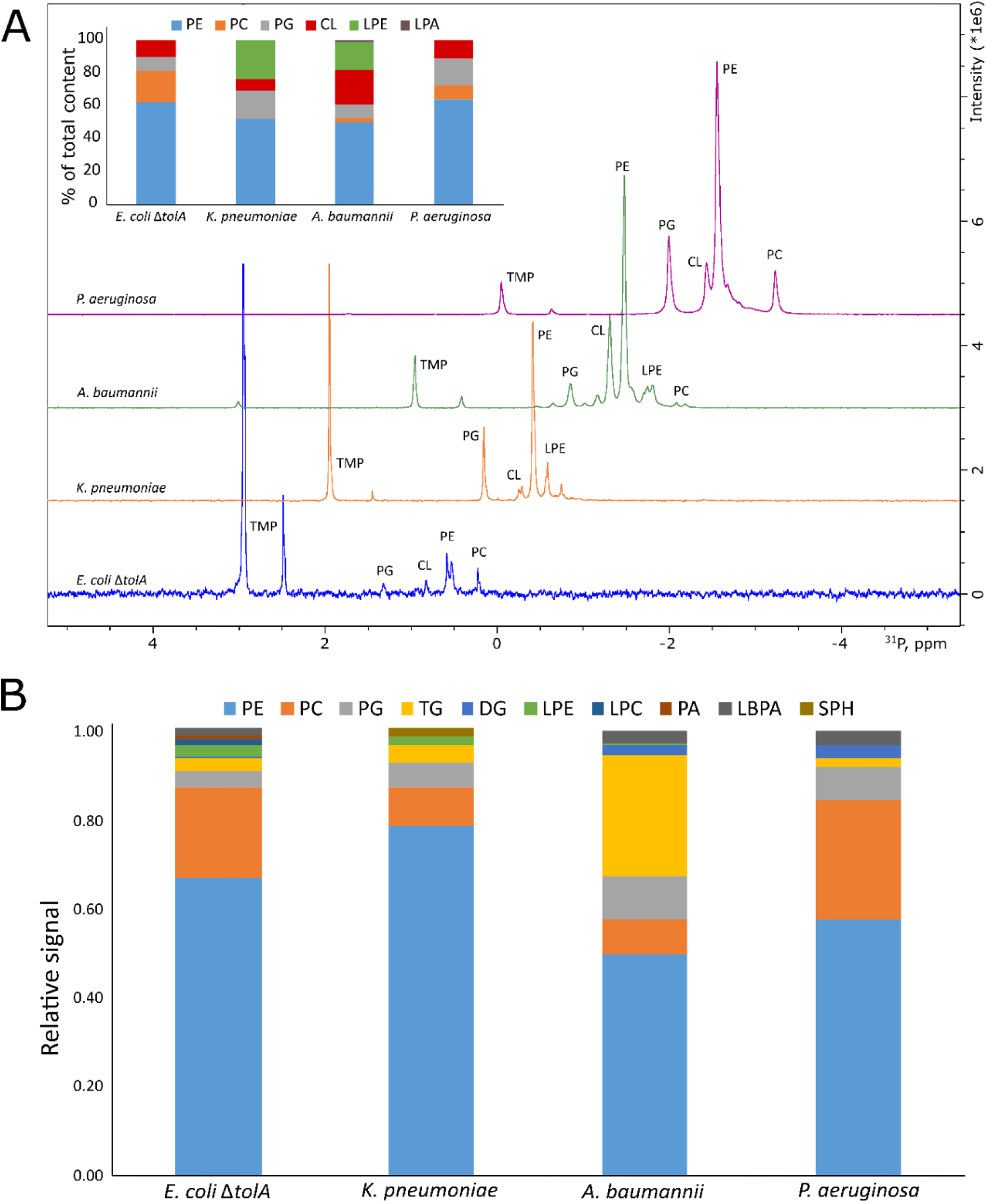
Summary of lipidomics analysis of bacterial lipid isolates by 31P NMR (A) and Mass Spectrometry (B). Intersection in the panel A shows quantified composition of the lipids in %. Traces of the NMR spectra were shifted up and right to ease the display. Here PE – phosphatidylethanolamine; PC – phosphatidylcholine; PG – phosphatidylglycerol; TG – triglycerides; DG – diglycerides; LPE – lyso-phosphatidylethanolamine; LPC – lyso-phosphatidylcholine; CL – cardiolipin; PA – phosphatidic acid; LBPA – lysobisphosphatidic acid; SPH – sphingomyelin.

The results from the MS analysis (Figure 3B) revealed that the lipid isolates, in addition to the species detectable by ^31^P NMR, also contained triglycerides (TG), diglycerides (DG) and various minor lipids. Although our MS approach only allowed for relative quantification within each sample, the results are in qualitative agreement with the ^31^P NMR in that PE was the dominant lipid class in all samples. The relative abundance of PC concurs with ^31^P NMR indicating that PC is the highest for *E. coli* Δ*tolA* and *P. aeruginosa*. While PC is commonly found in *P. aeruginosa* this phospholipid is normally not present in the bacterial membrane of *E. coli* strains (43, 46). However, the lipid composition is known to be dependent on the environment they are being cultured in and the rich lysogeny broth (LB) medium used in this work is most probably the source of PC. MS further allowed the detection of lipids not containing phosphorous, and the results show a notable TG signal for *A. baumannii*, but not for the other samples. Traces of lysobisphosphatidic acid (LBPA) and sphingomyelin (SPH) were detected for *K. pneumoniae*, likely acquired from the LB growth media as these lipids are not expected in bacteria. Sphingolipids, however, have been detected for *K. pneumoniae* before (8), and this class of lipids plays a role in establishing the balance between host and pathogen upon infection (47). Generally, bacteria are missing biosynthetic pathways for atypical lipids, but they are able to utilize fatty acids, phospholipids or their precursors available in the environment to be able to adapt and, for example, to evade the host immune defense system (7, 47). In the current study, a rich LB medium based on yeast extracts was used, which could potentially be the source of atypical lipids or their precursors (48). Overall, the results demonstrated that *K. pneumoniae* has a more diverse lipidome of minor lipids.

In general, quantitative comparison between NMR (quantitative relative abundance) and MS (non-quantitative relative abundance) were interpreted conservatively. The higher sensitivity of MS together with not being restricted to only observing ^31^P containing lipids however allowed us to compare the MS “fingerprint” of the total lipid isolates to the “fingerprint” of the OMV lipids produced by the same bacteria.

The comparison between the lipid isolate samples and the corresponding OMVs (Figure 4) provided a qualitatively good match between the lipid compositions of the OMVs and the isolates from the whole bacteria. The results showed that PE is the dominating lipid, and over all the same minor lipids are identified. The main deviation was observed for *A. baumannii* and *E. coli* Δ*tolA*, who showed different distributions of PC between the outer membrane vesicles and the whole cell isolate. *A. baumannii* had more of PC in the outer membrane and *E. coli* Δ*tolA* had less PC in the outer membrane (Figure 4). TG is utilized by bacteria as the energy storage lipid and thus, higher levels of TG in the whole cell isolate would be expected to originate from the interior of the bacteria (46, 49). In the case of the Δ*tolA* mutation in *E. coli*, the Tol-Pal membrane maintenance system is affected, which might potentially affect the degree of TG incorporation into the outer membrane (29).

**Figure 4.**
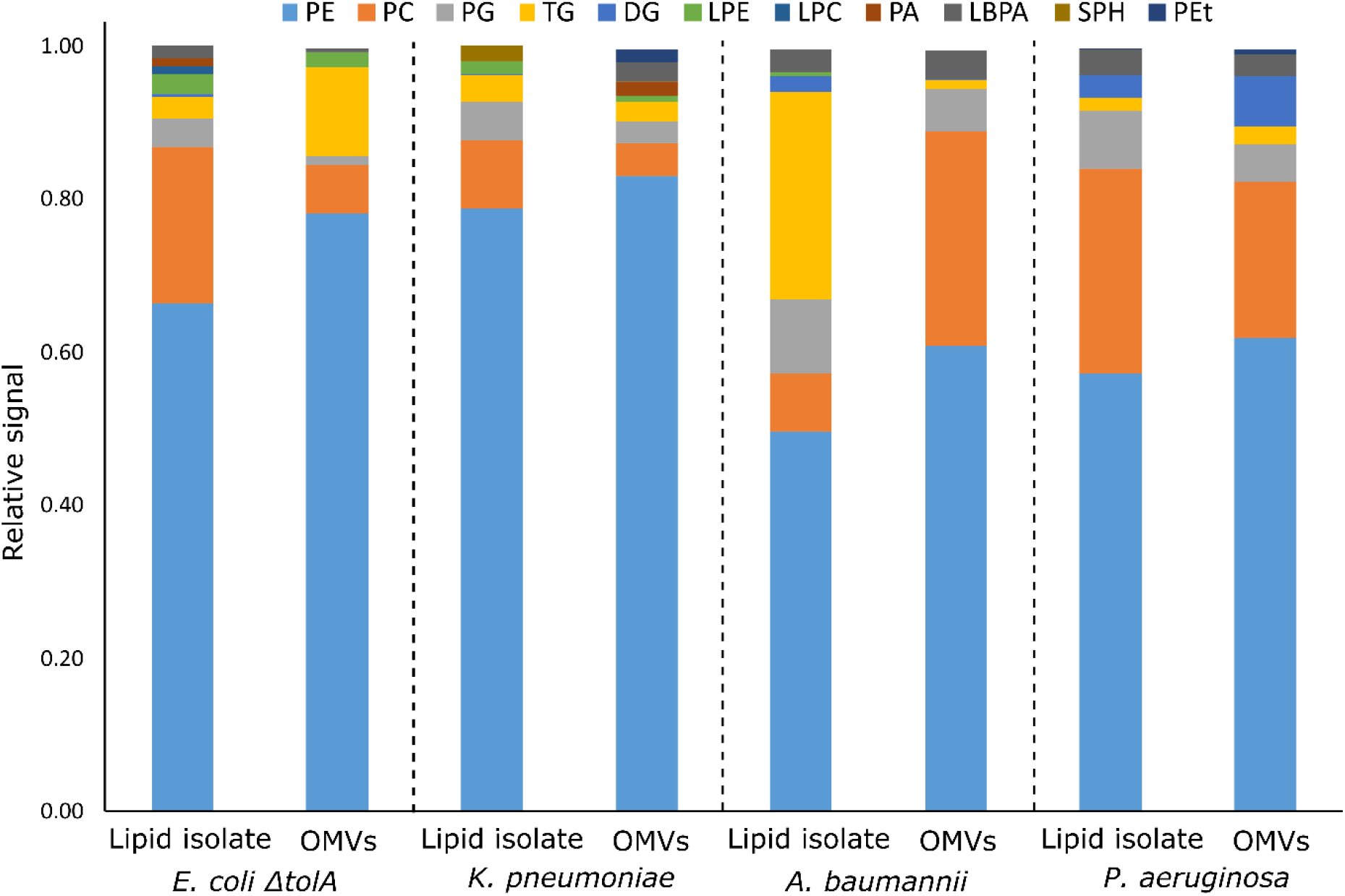
Mass Spectrometry fingerprints of the lipid compositions of the OMVs in comparison to the lipid isolates of corresponding strains: *E. coli* Δ*tolA, K. pneumoniae, A. baumannii* and *P. aeruginosa*. Here PE – phosphatidylethanolamine; PC – phosphatidylcholine; PG – phosphatidylglycerol; TG – triglycerides; DG – diglycerides; LPE – lyso-phosphatidylethanolamine; LPC – lyso-phosphatidylcholine; SPH – sphingomyelin; PA – phosphatidic acid; LBPA – lysobisphosphatidic acid; PEt – phosphatidylethanol.

The other notable deviation is for *K. pneumoniae*, where the background level is notably more diverse than for other samples and the outer membrane had more hits for LBPA, PA and PEt (phosphatidyl ethanol), but had no signal for SPH. The lipid isolate of *E. coli* Δ*tolA* also demonstrated higher diversity of background lipids that were not reflected for OMVs (Figure 4). This could possibly be attributed to the same mutation in the membrane maintenance system (29).

The MS approach used also allowed the analysis of the fatty acid composition of the detected lipid species. The analysis was limited to the PE phospholipid as it is the most dominant phospholipid in all samples. Out of the detected fatty acids, the hits with a relative abundance higher than 1 % were selected for analysis (Table S2). Mono-unsaturated fatty acids dominated in all samples. The top hits with relative abundance above 10 % are mono-unsaturated or saturated fatty acids with a chain length of 16 or 18 carbon atoms in different combinations. The results also suggested higher preferences of *K. pneumoniae* for shorter fatty acid chains with 14 carbons, or chains with odd number of carbons (17 or 19) instead of 18 carbons. For the *E. coli* Δ*tolA* sample, a fatty acid with 20 carbons and 4 double bonds was detected. When comparing the fatty acid composition of the OMVs to the lipid isolate from the whole bacteria it was observed that *E. coli* Δ*tolA* had similar lipid compositions for the OMVs and the native cell, while for the ESKAPE strains some differences could be observed. In particular this was evident for *K. pneumoniae* where the OMVs to a higher extent contained even numbered fatty acid chains (e.g., PE 16:0_16:1) compared to the whole bacteria where the PE 16:0_17:1 and PE 16:0_19:1 were the most prominent. This could affect its lipid phase and physical properties like permeability, flexibility, fluidity. The dynamic nature of membrane lipid metabolism is another factor that can allow bacteria to survive antimicrobials by modifying lipids, fatty acids distribution or their whole lipid composition (45, 50-54).

In conclusion, the OMVs are shown to inherit the lipid composition of the parent bacterial membrane, including strain specific variations. Thus, the OMVs showed potential to be an elegant model system for targeted studies, carrying authentic bacterial outer membrane in a highly stable and reasonably sized nano-construct.

### Interaction studies by SPR

The potential to use OMVs as realistic models for the bacterial membrane in interaction studies was explored. As proof-of-principle, the interactions between OMVs and AMPs were measured using SPR.

For the SPR analysis, samples of OMVs or liposomes were immobilized at the surface of L1 chips, and the dilution series of the antimicrobial peptides were run over the layer formed by the OMVs, measuring the response. A set of synthetic cyclic antimicrobial peptides (AMPs) with alternating- or clumped distribution of charged (Lysine or Arginine) and hydrophobic (Tryptophan) amino acids was used: c-WKWKWK, c-WRWRWR, c-WWWKKK and c-WWWRRR (Figure 5). The set was selected as a useful chemical model to study the effects of charge distribution on the interactions. The peptide c-LWwNKr (Figure 5) was included in the study as a control that is chemically similar, but has no antimicrobial activity (MIC > 256 μg/ml) (37-42)

**Figure 5.**
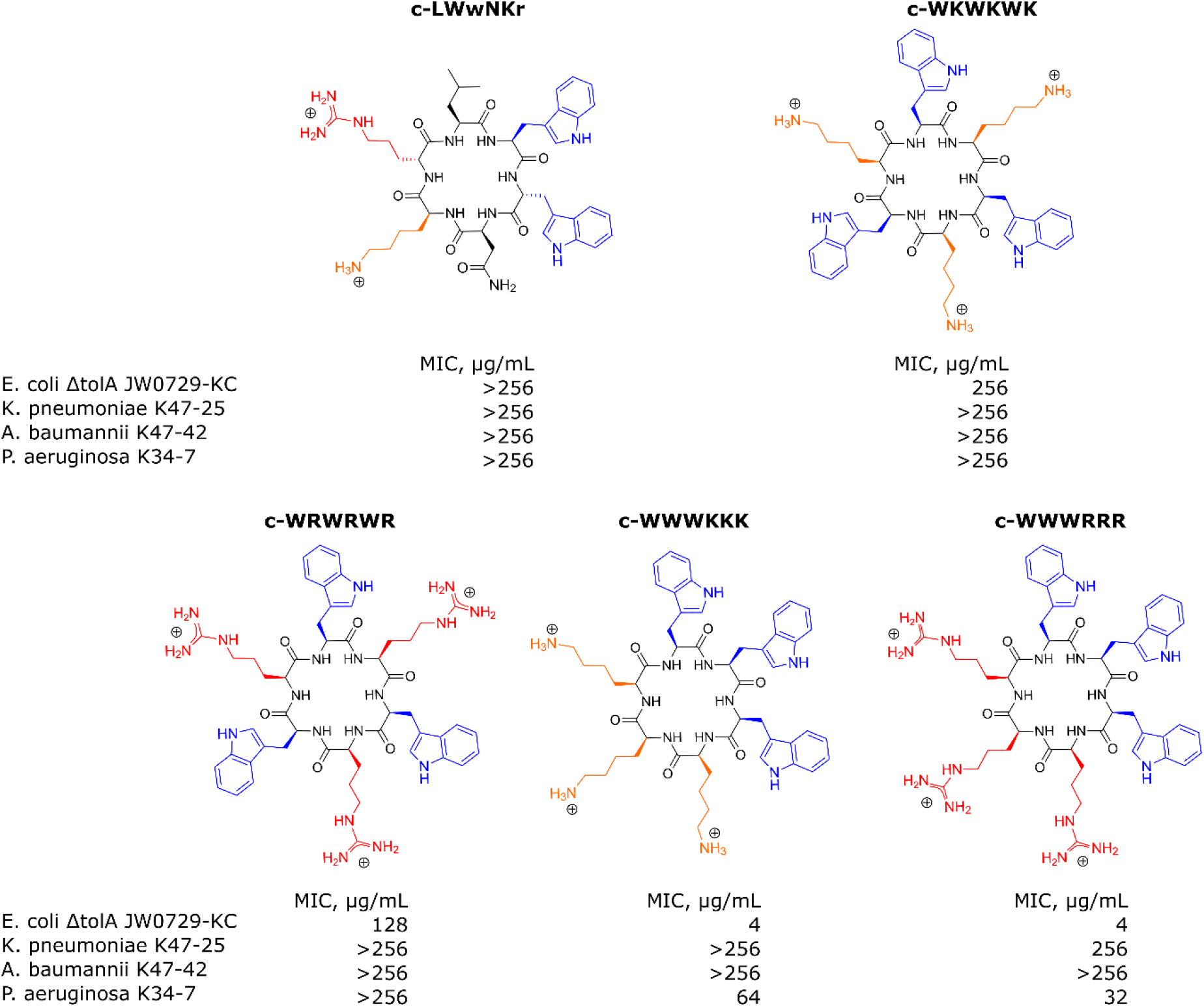
Structures of antimicrobial peptides used in this study and their minimal inhibitory concentrations (MIC) values determined for *E. coli* Δ*tolA* JW0729-KC, *K. pneumoniae* K47-25, *A. baumannii* K47-42 and *P. aeruginosa* K34-7. Hydrophobic Tryptophans highlighted in blue, positively charged Arginines and Lysines are highlighted in red and orange correspondingly.

To analyze the interaction process the model described by Figueira et al., 2017 was used (55). With this model, the dissociation constant K_D_ and the disassociation rate k_off_ were obtained. These constants allowed us to calculate k_on_ rate, using equation 4. Figures 6A-D present examples of the raw response curves obtained during a run and the analytical processing steps of data fitting and analysis. The results for all samples are summarized in Figures 6E-G, and Tables S3, S4 and S5 in Supplementary materials.

**Figure 6.**
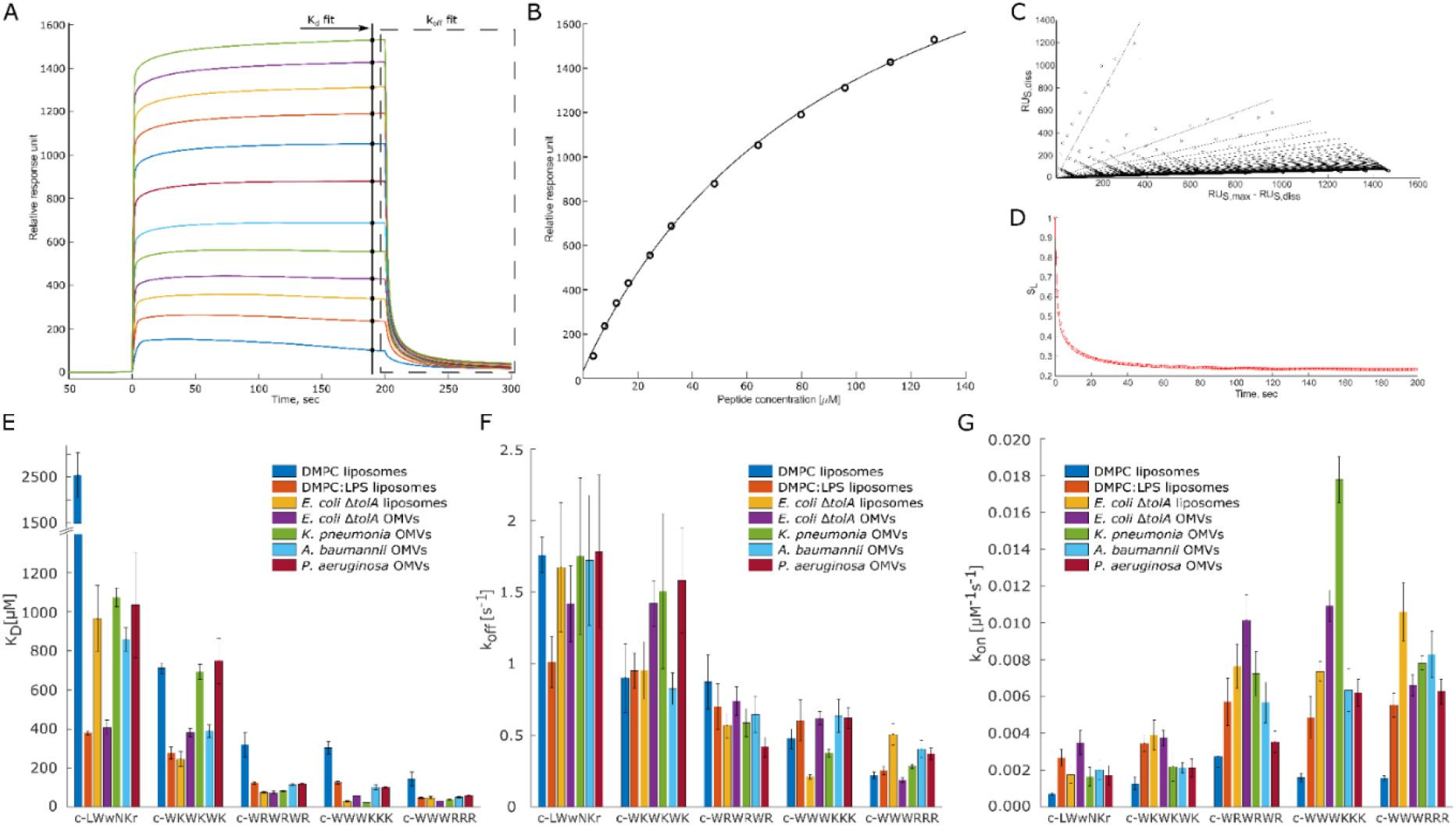
Overview panel of the SPR method and results. **A** – example of typical raw response signals collected during a run. Marked data points used for fitting to models to obtain K_D_ panel **B**) and k_off_ values. To obtain k_off_ values the raw data of the marked region first was linearized (panel **C**) and then a model was fitted into in-time changing linear coefficients (panel **D**). Panel B, C and D represent typical data fit plots for our work. Panels **E** and **F** summarize measured K_D_ and k_off_ values. Panel **G** shows calculated k_on_ values. Three parallels were performed for each combination. Values are available in Table S3, S4 and S5 in supplementary materials. T-test statistical analysis of the data is summarized in Table S6 in supplementary materials. Results for DMPC and DMPC:LPS (90:10) are reproduced from Jakubec et al. (40).

The minimum inhibitory concentration (MIC) of the peptides against the bacterial strains used in this study was determined (Figure 5). The non-resistant *E. coli* Δ*tolA* displayed strong binding (low K_D_) and high antimicrobial activity (MIC = 4 μg/mL for both c-WWWKKK and c-WWWRRR). However, for the resistant strains none of the peptides showed any activity (MIC ≥ 256 μg/mL), except c-WWWKKK and c-WWWRRR that were inhibiting growth of *P. aeruginosa* at ≥64 μg/mL and ≥32 μg/mL, respectively. This contrasts with the fact that 3 out of the 4 antimicrobially active peptides bind as efficiently to the surface of the OMVs from resistant strains as to the OMVs from susceptible strains. The c-WKWKWK displayed a weaker interaction with the OMVs from *K. pneumoniae* and *P. aeruginosa* and the control peptide c-LWwNKr showed no antimicrobial activity in all cases as expected.

### Quantification of the non-lipid components’ contribution

Comparing interactions of the peptides with *E. coli* Δ*tolA* OMVs and liposomes assembled from the lipid isolate of the same strain gives an indication of how the kinetic barrier imposed by LPS together with membrane proteins and other native components affects the interaction. For lysine-containing peptides, both the k_off_ and k_on_ rates were positively affected for the interactions with the OMVs, but the effect was more pronounced for k_off_, resulting in an overall increase in K_D_ A corresponding increase in the k_on_ and k_off_ rates for the lysine-containing peptides was also observed in the recently published comparison between homogeneous DMPC liposomes and DMPC liposomes containing 10 % LPS (40) (results included in Figure 6E-G). In this case the balance was however different, resulting in slightly improved K_D_ and K_p_.

The opposite picture is observed for the c-WWWRRR peptide, where the introduction of the LPS barrier and other components decreased both the k_off_ and k_on_ rates, shifting the balance towards decreased dissociation. In the recent study with DMPC liposomes with and without LPS, a similarly improved K_D_ was also observed, but in that case, it was driven by a significant increase of the k_on_ rate (40). This peptide demonstrated the strongest binding of all tested peptides, and also had the lowest MIC value for *E. coli* Δ*tolA*. For the control peptide, the native OMV mostly contributed to an increase in association rate, k_on_, and a slight decrease in dissociation rate. The resulting change in the K_D_ for this sample was the most prominent among the studied peptides. This observation was fully consistent with the behavior in the DMPC/DMPC+LPS system, suggesting that LPS is likely responsible for the stronger binding of c-WWWRRR.

### Interactions of OMVs produced by ESKAPE strains

The overarching focus was to isolate and characterize OMVs from resistant clinical isolates of ESKAPE pathogens, and to explore their applicability in interaction studies with antimicrobial model compounds. Comparison of the results from the interaction studies with OMVs to those with liposomes made from *E. coli l*ipid isolate showed that the trend is consistent, with the clumped peptides being the strongest binders and the control peptide and c-WKWKWK being the weakest, reflected by both K_D_ and k_off_ (Figure 6 E-G). There are, however, notable peculiarities in the results. The c-WWWKKK peptide was the peptide in this study that had the highest affinity for OMVs from *K. pneumoniae* and was even binding stronger to the OMVs than to the *E. coli* lipid isolate liposomes. The OMVs from *K. pneumoniae* had a prominent positive effect on the k_on_ rate and only a slight increase in the k_off_ rate, resulting in an approximately 5-fold lower K_D_ compared to the K_D_ for other resistant strains. The outer membrane of *A. baumannii* and *P. aeruginosa* had on the other hand a negative effect on the k_off_ rate and K_D_ for this peptide.

The results for the control peptide, c-LWwNKr, displayed a strong preference for the outer membrane of *E. coli* Δ*tolA* over the lipid isolate liposomes. A similar increase was observed in DMPC+LPS liposomes compared to pure DMPC liposomes, and in both cases this increase is driven by an increase in k_on_. Even though this peptide is inactive, it exhibits this apparent affinity for LPS. This increase was not observed in the resistant strains. However, similar trends in interaction of resistant strains and the c-WKWKWK peptide could be found. Here the k_on_ rate (Figure 6E) was reduced similarly for OMVs from the resistant strains compared to the *E. coli* lipid isolate liposomes, while for the OMVs from the non-resistant *E. coli* Δ*tolA* there was no statistically significant change in k_on_. The k_off_ rate (Figure 6F) was slightly decreased for *A. baumannii* while significantly increasing for the other strains, including the susceptible *E. coli* Δ*tolA*. Thus, as a result the K_D_ values (Figure 6G) of OMVs from *A. baumannii* is similar with OMVs from *E. coli* Δ*tolA*, while for OMVs from *K. pneumoniae* and *P. aeruginosa* a twofold increase in K_D_ was displayed.

For the interactions with the c-WRWRWR peptide, the OMVs from the resistant strains displayed a decrease in k_on_ rates compared to the *E. coli* lipid isolate liposomes, especially for *A. baumannii and P. aeruginosa*. For the susceptible *E. coli* Δ*tolA*, the k_on_ rate increased for this peptide compared to the lipid isolate liposomes. The c-WWWRRR peptide showed similar behavior for all samples, slowing down both k_on_ and k_off_ rates, only slightly changing the balance of K_D_.

The overall trend when comparing the peptides to each other is, however, that the peptides that display a strong affinity for the *E. coli* lipid isolate and the OMVs of the susceptible strain, also display a strong affinity for the OMVs from the resistant strains. The only exception is the c-WKWKWK peptide with a moderate affinity, which displayed a weaker affinity for two out of the three resistant strains.

OMVs from ESKAPE pathogens further appear to have inherited the strain specific properties of the outer membranes, reflecting the native surface of the bacteria. The different properties of the vesicles originating from different strains resulted in observed differences in their interactions with AMPs measured by SPR. Further, comparison to liposomes made from the whole bacteria lipid isolate allowed us to distinguish between the contributions from the lipids and the other outer membrane components, presumably dominated by LPS. The results demonstrated that the affinity of the AMPs to ESKAPE strains did not correlate with their activity. The strongest binder was inactive towards the resistant strains, while being active in non-resistant strain with comparable binding parameters. Some potential explanations why there are no close correlations between the binding affinity and the antimicrobial effect could be that the resistant bacteria either maintain the membrane integrity upon association, or the bacteria had a mechanism to efficiently deactivate the AMPs (27, 28). Another possible explanation why the peptides would bind but did not cause bacteriostasis is that the peptide did not manage to penetrate the kinetic conditions set by LPS and its modifications (6, 9, 10).

## Conclusions

The traditionally used synthetic membrane models for the bacterial membrane cannot effectively reconstruct the full complexity of native bacterial membranes. Herein, we have for the first time shown that the bacteria can do the hard work for us and produce sufficiently large quantities of native OMVs that enable interaction studies by SPR between compounds of interest and authentic bacterial outer membrane. The isolated OMVs were found to inherit the strain specific variations of the outer membranes composition, which in turn dictated the kinetic rates of the interaction processes.

Thus, we have provided proof of principle that native OMVs produced by clinical isolates of Gram-negative ESKAPE strains, carrying specific antimicrobial resistance genes, can be used as tools to study processes on the bacterial surface. This sets the conditions to enable direct quantification of, for example, the effect of induction by antibiotics, or the response to host-pathogen interactions.

## Materials and methods

### Materials

All common chemicals and solvents were purchased from Sigma Aldrich (Merk KGaA, Darmstadt, Germany). *Escherichia coli* Δ*tolA* strain was obtained from National BioResource Project (NIG, Japan): E. coli (Keio Collection JW0729-KC). Strains of clinical isolates of multi-drug resistant *K. pneumoniae* K47-25, *A. baumannii* K47-42 *and P. aeruginosa* K34-7 strains were provided by The Norwegian National Advisory Unit on Detection of Antimicrobial Resistance (K-res), University Hospital of Northern-Norway (UNN). The peptides c-WKWKWK, c-WRWRWR, c-WWWKKK, c-WWWRRR and c-LWwNKr were synthesized as previously reported in (40, 42). PhosSTOP tablets were purchased from Roche via Sigma Aldrich. Lipids were purchased from Avanti Lipids via Sigma Aldrich.

### Minimum inhibitory concentration (MIC) determination

MIC for all test peptides was determined following EUCAST clinical standard (56). Briefly, 3-4 colonies were picked by a cotton swab from freshly plated cultures on agar and resuspended in sterile saline solution. Turbidity of inoculum was adjusted to 0.5 McFarland standard units prior to 100x dilution in Mueller-Hinton broth. Antimicrobial peptides were dissolved in the broth at the stock concentration of 512 μg/mL. MIC was determined in series of dilutions of antimicrobial peptides at concentrations ranging from 256 to 2 μg/mL in microdilution plates, keeping concentration of bacterial cells at constant. Positive control of bacterial culture diluted with broth and negative control of blank broth were included for each bacterial strain. The MICs were determined after incubation of the plates at 37 °C for 18 hours and red as the concentration of antimicrobial peptide at the next to the last well where bacterial growth was detected.

### Outer membrane vesicles isolation

Bacterial cultures were grown in lysogeny broth (LB) media at 37 °C at 120 rpm until stationary growth phase for 16 hours and harvested by centrifugation at 10,000 × g for 20 min. For further lipidomic analysis the bacterial pellet was lysed by immediate resuspension in 5 mL of TRIS buffer (10 mM TRIS, 100 mM NaCl and 5 mM KCl, pH 8.0) supplemented with 1 tablet of PhosSTOP, 2 mg/mL of 2-butoxyphenylboronic acid (BPBA), and 500 μL of mixture of mutanolysin (50 μg), lysozyme (10 mg), RNAase (200 μg) per 1 mL of glycerol/PBS 1:1 (the stock stored at -80 °C). The mixture was left overnight at room temperature on a soft shaker and then frozen at -80 °C before being freeze dried.

For isolation of OMVs the collected supernatant was filtered through 0.45 μm PES bottle-top vacuum filters and further upconcentrated to approx. 50-100 mL using Sartorius VivaFlow 200 30,000 PES membrane according to the instruction manual. Concentrate was filtered through 0.22 μm filters using a syringe, and then was ultracentrifuged (Beckman Coulter, Optima XPN-100) at 30,000 rpm for 2 hours. The supernatant was discarded, and the pellet was resuspended in TRIS buffer, following another run of centrifugation at 30,000 rpm for 2 hours. The remaining pellet, which contains the outer membrane vesicles, was finally resuspended in 300 μL TRIS buffer and inspected by negative-stain electron microscopy to assess the quality of the samples and analyzed on Zetasizer (see below). Protein concentration was used as a reference for the samples and was measured by Bradford protein assay from Bio-Rad.

### Lipids isolation

To isolate lipids a modified Bligh and Dyer method was used (57, 58). Briefly, lysed bacterial pellet was freeze dried for 48 hours after being treated with the enzymatic mixture as described above. The freeze-dried material was mixed with 10-15 mL of dichloromethane/methanol mixture (2:1), properly vortexed and centrifuged (6,000 × g for 10 min). The organic solution was decanted to a separation funnel and the remaining pellet was mixed with another 10-15 mL of the same dichloromethane/methanol mixture. After vortexing the dispersion was transferred to the same separation funnel. The same volume of 5 % NaCl solution in water was added to the funnel. The content of the funnel was gently mixed, and the mixture was left for the phase separation. After separation, the bottom organic phase was collected, and the remaining water phase was washed by addition of the same volume of dichloromethane/methanol mixture. Again, the mixture was allowed to form the phases and the bottom organic one was collected and combined with the previously collected fraction. The collected pooled organic mixture was filtered through filter paper and evaporated on a rotary evaporator. The resulting lipid film of *E. coli* Δ*tolA* was used to prepare liposomes for SPR analysis (by hydration with TRIS buffer, see below). Otherwise, the dried lipid films were redissolved in smaller volume of isopropanol and transferred to a smaller pre-weighted glass container through a cotton filter. After drying the lipid isolates under a nitrogen stream the resulting lipids were weighed. These isolates were used to prepare samples for ^31^P NMR analysis by dissolving in CUBO solvent or for MS analysis by dissolving in isopropanol (see below).

Lipid isolation from OMVs was performed in a similar fashion but with a simpler procedure. Freeze dried OMVs were resuspended in 3-5 mL of 2:1 mixture of dichloromethane and methanol, sonicated for 2 minutes of 2 sec on/8 seconds off cycles on ice using probe sonicator (Ultrasonic processor 500W, Sigma Aldrich, USA). 3-5 mL of 5 % NaCl was added to the samples and after thorough vortexing the phases were separated by centrifugation at 3,000 × g for 5 min. The organic solution was transferred to a clean glass vial with glass pipet. The remaining aqueous phase was mixed with another 3-5 mL of the same dichloromethane/methanol mixture, vortexed and centrifuged. After separation, the organic phases were pooled together into the glass vial and evaporated under nitrogen. Dried isolates were redissolved in a smaller volume of isopropanol and transferred into a smaller pre-weighted glass container through a cotton filter. After drying under nitrogen, the lipid isolates were weighed. The samples were dissolved in isopropanol for mass spectrometry analysis.

### Mass Spectrometry

All lipid samples were dissolved in isopropanol at 1 mg/mL concentration. Identification and relative quantification of lipids were done on a Thermo IdX Tribrid Orbitrap MS with a Thermo Vanquish UPLC. Separation was done on a Waters Acquity Premier BEH C18 1.7 um 2.1 x 100 mm column with gradient elution. Mobile phase A consisted of 50 % ACN in MilliQ-water and mobile phase B of 47.5 % ACN, 47.5 % isopropanol and 5 % MilliQ-water. Both mobile phases were buffered with 1 mM ammonium formate and added formic acid to pH 3. The gradient started with 30 % B, followed by a linear increase to 75 % B at 20 minutes, again followed by a linear increase to 95 % B at 25 minutes, which were held for 5 minutes, giving a total analysis time of 30 minutes. The injection volume was 2 μl, the column temperature 60 °C, the flow rate 0.6 ml/min, and the sample compartment was held at room temperature to avoid precipitation of lipids.

Identification of lipids were done by MS^2^ and MS^3^ experiments with the orbitrap mass analyzer. The analysis was done with positive ionization, the orbitrap full scan analysis was run with a resolution of 120,000, while MS^2^ and MS^3^ experiments were run at 15,000. Identification was obtained by first running a full scan in the mass range m/z 250-1500, followed by data dependent MS^2^ (ddMS^2^) where the most intense ions were selected for HCD fragmentation with stepped collision energy (25, 30, 35 %). Based on the results from the first fragmentation the ions could be sent for a second MS^2^ analysis if the result showed that the molecule was a phosphocholine (based on m/z 184 corresponding to the head group) for full identification by CID fragmentation. If the first ddMS^2^ showed that the molecule was a triacylglycerol (by neutral loss of a fatty acid) the ions were sent for MS^3^ analysis by CID fragmentation for full identification. All other lipids were identified after the first ddMS^2^ fragmentation. To identify as many lipids as possible in the samples 5 consecutive injections of a pooled sample were analysed with MS^n^ fragmentation. In the first injection the most intense signals are fragmented for identification, thereafter, put on an exclusion list for injection 2 so that signals with lower intensity are fragmented and identified in the second injection. The process is repeated for all 5 injections to ensure identification of all lipids with significant signal intensity. The samples were analyzed in full scan with a resolution of 120,000, and after analysis the results were aligned with the results from the MS^2^ and MS^3^ experiment for identification. Lipid identification and relative quantification were done with the software LipidSearch from Thermo. More details can be found in S2_MS_Method.pdf supplementary material.

### Liposomes preparation

The lipid film from *E. coli* Δ*tolA* culture lipid isolate was prepared by removing the solvent under vacuo at 45 °C on a rotavapor with final step of holding at vacuum of 55 hPa at room temperature for 3 hours. The lipid film was then hydrated with TRIS buffer. Before extrusion through 100 nm polycarbonate membrane filters (Millipore) the liposome dispersions were subjected to 5 min of sonication cycles (5 sec impulse followed by 10 sec hold) at room temperature using probe sonicator. Size and zeta-potential of resulting liposomes were analyzed by zetasizer (see below).

### Size and zeta potential measurements

Samples of isolated OMVs and freshly extruded liposomes prepared from lipid isolate dispersed in TRIS buffer were subjected to size and zeta potentials measurements using a Malvern Zetasizer Nano ZS (Malvern, Oxford, UK) instrument. In addition to size measurements, the polydispersity index (PDI) was estimated to assess population homogeneity. For comparison zeta-potential of the bacterial cells of the presented strains were determined by growing a 10 mL mini-prep of bacterial culture in LB media for 6 hours and measuring the zeta-potential by diluting the samples in TRIS buffer. All zeta-potential measurements were performed using the disposable folded capillary cells (DTS1070). All samples were measured in triplicates at 25 °C.

### Electron microscopy

For negative stain electron microscopy freshly carbon-coated 400 mesh grids were used. The grids were absorbed on the droplets of isolated OMVs for 5 min, washed with double distilled water and stained with 1:9 mixture of 3 % uranyl acetate and 2 % methylcellulose for 2 min. The grids with the sample were picked up with loops, excess uranyl acetate was removed with filter paper, and the loops with the grids were dried on the loop holder for 10 min prior the imaging. The micrographs were obtained with HT7800 Series transmission electron microscope (Hitachi High-Tech Corp., Tokyo, Japan) operating at accelerated voltage of 100 kV coupled with Morada camera.

### SPR assays

OMVs isolates resuspended in TRIS buffer at 0.5 mg/mL protein concentration were used as is without additional treatment. It was not possible to keep track on the final concentration of liposomes prepared from lipid isolate. For this sample optimal dilution was empirically determined by monitoring coverage of the chip. The SPR experiments were performed as was described in (40, 42). Briefly, the analysis was carried out with a T200 Biacore instrument (GE Healthcare, Chicago, Ill, USA), at 25 °C. Samples of OMVs or liposomes were immobilized on the surface of L1 chip (Cytiva, Marlborough, MA, USA) at 2 μL/min flow rate for 2400 sec. Chip coverage was assessed by injection of 0.5 mg/ml of BSA at 30 μL/min flow rate for 60 sec. Change in response units (RU) of less than 400 was considered as sufficient coverage of the chip. The peptides were tested at increasing range of concentrations (from 4 to 128 μM for c-WRWRWR, c-WWWKKK and c-WWWRRR or 24 to 768 μM for c-WKWKWK and c-LWwNKr) in 10 mM HEPES buffer (with 150 mM NaCl). The peptides were injected over immobilized samples at 15 μL/min flow rate for 200 sec followed by 400 sec of dissociation phase. After each injection, the surface of immobilized vesicles was stabilized by three subsequent injections of 10 mM NaOH at 30 μL/min flow rate for 30 sec each. Each run was concluded with cleaning of the chip surface by subsequently injecting 10 mM CHAPS, 40 mM octyl-β-D-glucopyranoside and 30 % ethanol for 1 minute at 30 μL/min flow rate. The control channel of the chip was treated the same way, except only HEPES buffer was injected instead of peptide solutions. The resulting data sets were processed and analyzed by in-house developed MATLAB scripts (MATLAB R2020a; available at: https://github.com/MarJakubec). K_D_ was obtained using steady-state fit (equation 1) and k_off_ rate constant was determined using formalism suggested by Figuera *et al*. 2017 (equations 2 and 3). The association k_on_ rate was calculated from obtained K_D_ and k_off_ values using equation 4.4.

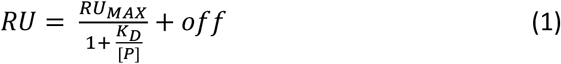

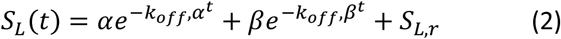

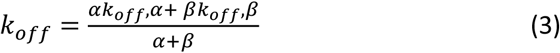

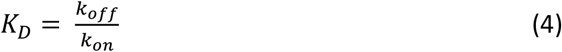

where RU is measured SPR response; RU_MAX_ is RU response at saturation; [P] is peptide concentration, and *off* is the offset of the response; S_L_ is the linearised ratio of responses of the solute, which is plotted against the time of dissociation; α and β are individual populations, and S_L,r_ is the retained solute fraction.

### ^31^P NMR Spectroscopy

NMR spectra were acquired using a Bruker 400 MHz Avance III HD spectrometer equipped with an RT SmartProbe. Lipid isolates from bacterial membranes were dissolved in 500 μL of CUBO solvent (800 mM guanidine chloride dissolved in 3:1 mixture of dimethylformamide (DMF) and trimethylamine). For lipid quantification trimethyl phosphate (TMP) standard was added to the samples at the concentration of 0.7 μM. All experiments were acquired at 298 K in 5 mm NMR tubes. Spectra were processed in TopSpin 4.1.4 (Bruker, Billerica, MA, USA).

## Supporting information

Supplementary_MSmethod_OMVs as bacterial membrane models

Supplementary_tablesandfigure_OMVs as bacterial membrane models

## Abbreviations

OMVs: outer membrane vesicles
AMPs: antimicrobial peptides
SPR: surface plasmon resonance
PLs: phospholipids
LPS: lipopolysaccharides
NMR: nuclear magnetic resonance spectroscopy
MS: mass spectrometry

## Acknowledgement

The authors acknowledge research funding support from CANS Centre for New Antibacterial Strategies (TFS grant No. 18_CANS_AS) at UIT The Arctic University of Norway, the Digital Life project DigiBiotics (ID 269425) granted by the Research Council of Norway as well as funding from NordForsk for the Nordic University Hub project #85352 (Nordic POP, Patient Oriented Products). The APC was covered by the open access publishing fund, UiT. Prof. Hanne Cecilie Winther-Larsen and Verena Mertes at Department of Pharmacy, University of Oslo, Norway, are greatly acknowledge for hosting a mobility action, scientific discussions and knowledge transfer on OMVs production. TEM-imaging was done with help from The Advanced Microscopy Core Facility, Department of Medical Biology, UiT The Arctic University of Norway and the peptides acquired from the Digibiotics pipeline.

## Author contributions

Conceptualization: MB. Sample preparation: MB, VS. Methods optimization: MB, MJ. SPR analysis: MB, VS, MJ. Lipidomics: MB, TV, MJ. Synthesis: TK. Data analysis: MB, VS, MJ. Original draft: MB. Draft revision: MB, JC, JI, GF, JE. Project design and funding acquisition: GF, JI, JP, and JE. All authors read and commented on the final version of the manuscript.

## REFERENCES

1. WHO. Antimicrobial resistance: global report on surveillance: World Health Organization; 2014.

2. Antimicrobial_Resistance_Collaborators. Global burden of bacterial antimicrobial resistance in 2019: a systematic analysis. Lancet. 2022;399(10325):629–55.

3. Mulani MS, Kamble EE, Kumkar SN, Tawre MS, Pardesi KR. Emerging Strategies to Combat ESKAPE Pathogens in the Era of Antimicrobial Resistance: A Review. Front Microbiol. 2019;10:539.

4. WHO. WHO publishes list of bacteria for which new antibiotics are urgently needed 2017 [Available from: https://www.who.int/news/item/27-02-2017-who-publishes-list-of-bacteria-for-which-new-antibiotics-are-urgently-needed.

5. Delcour AH. Outer membrane permeability and antibiotic resistance. Biochim Biophys Acta. 2009;1794(5):808–16.

6. Llobet E, Martínez-Moliner V, Moranta D, Dahlström KM, Regueiro V, Tomás A, et al. Deciphering tissue-induced Klebsiella pneumoniae lipid A structure. Proc Natl Acad Sci U S A. 2015;112(46):E6369–78.

7. Hobby CR, Herndon JL, Morrow CA, Peters RE, Symes SJK, Giles DK. Exogenous fatty acids alter phospholipid composition, membrane permeability, capacity for biofilm formation, and antimicrobial peptide susceptibility in Klebsiella pneumoniae. Microbiologyopen. 2019;8(2):e00635.

8. Jasim R, Han ML, Zhu Y, Hu X, Hussein MH, Lin YW, et al. Lipidomic Analysis of the Outer Membrane Vesicles from Paired Polymyxin-Susceptible and -Resistant. Int J Mol Sci. 2018;19(8).

9. Olaitan AO, Morand S, Rolain JM. Mechanisms of polymyxin resistance: acquired and intrinsic resistance in bacteria. Front Microbiol. 2014;5:643.

10. Jeannot K, Bolard A, Plésiat P. Resistance to polymyxins in Gram-negative organisms. Int J Antimicrob Agents. 2017;49(5):526–35.

11. Beveridge TJ. Structures of gram-negative cell walls and their derived membrane vesicles. J Bacteriol. 1999;181(16):4725–33.

12. Roier S, Zingl FG, Cakar F, Durakovic S, Kohl P, Eichmann TO, et al. A novel mechanism for the biogenesis of outer membrane vesicles in Gram-negative bacteria. Nat Commun. 2016;7:10515.

13. Silhavy TJ, Kahne D, Walker S. The bacterial cell envelope. Cold Spring Harb Perspect Biol. 2010;2(5):a000414.

14. Akbarzadeh A, Rezaei-Sadabady R, Davaran S, Joo SW, Zarghami N, Hanifehpour Y, et al. Liposome: classification, preparation, and applications. Nanoscale Res Lett. 2013;8(1):102.

15. Sezgin E, Schwille P. Model membrane platforms to study protein-membrane interactions. Mol Membr Biol. 2012;29(5):144–54.

16. Bailey-Hytholt CM, LaMastro V, Shukla A. Assembly of Cell Mimicking Supported and Suspended Lipid Bilayer Models for the Study of Molecular Interactions. J Vis Exp. 2021(174).

17. Jackman JA, Cho NJ. Supported Lipid Bilayer Formation: Beyond Vesicle Fusion. Langmuir. 2020;36(6):1387–400.

18. Denisov IG, Schuler MA, Sligar SG. Nanodiscs as a New Tool to Examine Lipid-Protein Interactions. Methods Mol Biol. 2019;2003:645–71.

19. Scheidelaar S, Koorengevel MC, Pardo JD, Meeldijk JD, Breukink E, Killian JA. Molecular model for the solubilization of membranes into nanodisks by styrene maleic Acid copolymers. Biophys J. 2015;108(2):279–90.

20. Denisov IG, Grinkova YV, Lazarides AA, Sligar SG. Directed self-assembly of monodisperse phospholipid bilayer Nanodiscs with controlled size. J Am Chem Soc. 2004;126(11):3477–87.

21. Pirc K, Ulrih NP. α-Synuclein interactions with phospholipid model membranes: Key roles for electrostatic interactions and lipid-bilayer structure. Biochim Biophys Acta. 2015;1848(10 Pt A):2002–12.

22. Prenner EJ, Lewis RN, McElhaney RN. The interaction of the antimicrobial peptide gramicidin S with lipid bilayer model and biological membranes. Biochim Biophys Acta. 1999;1462(1-2):201–21.

23. Jakubec M, Bariås E, Furse S, Govasli ML, George V, Turcu D, et al. Cholesterol-containing lipid nanodiscs promote an α-synuclein binding mode that accelerates oligomerization. FEBS J. 2021;288(6):1887–905.

24. Elderdfi M, Sikorski AF. Interaction of membrane palmitoylated protein-1 with model lipid membranes. Gen Physiol Biophys. 2018;37(6):603–17.

25. Paulowski L, Donoghue A, Nehls C, Groth S, Koistinen M, Hagge SO, et al. The Beauty of Asymmetric Membranes: Reconstitution of the Outer Membrane of Gram-Negative Bacteria. Front Cell Dev Biol. 2020;8:586.

26. Haurat MF, Elhenawy W, Feldman MF. Prokaryotic membrane vesicles: new insights on biogenesis and biological roles. Biol Chem. 2015;396(2):95–109.

27. Balhuizen MD, van Dijk A, Jansen JWA, van de Lest CHA, Veldhuizen EJA, Haagsman HP. Outer Membrane Vesicles Protect Gram-Negative Bacteria against Host Defense Peptides. mSphere. 2021;6(4):e0052321.

28. Manning AJ, Kuehn MJ. Contribution of bacterial outer membrane vesicles to innate bacterial defense. BMC Microbiol. 2011;11:258.

29. Reimer SL, Beniac DR, Hiebert SL, Booth TF, Chong PM, Westmacott GR, et al. Comparative Analysis of Outer Membrane Vesicle Isolation Methods With an. Front Microbiol. 2021;12:628801.

30. Krüger M, Richter P, Strauch SM, Nasir A, Burkovski A, Antunes CA, et al. What an Escherichia coli Mutant Can Teach Us About the Antibacterial Effect of Chlorophyllin. Microorganisms. 2019;7(2).

31. Ruiz N, Falcone B, Kahne D, Silhavy TJ. Chemical conditionality: a genetic strategy to probe organelle assembly. Cell. 2005;121(2):307–17.

32. Paulsen MH, Engqvist M, Ausbacher D, Anderssen T, Langer MK, Haug T, et al. Amphipathic Barbiturates as Mimics of Antimicrobial Peptides and the Marine Natural Products Eusynstyelamides with Activity against Multi-resistant Clinical Isolates. J Med Chem. 2021;64(15):11395–417.

33. Ageitos JM, Sánchez-Pérez A, Calo-Mata P, Villa TG. Antimicrobial peptides (AMPs): Ancient compounds that represent novel weapons in the fight against bacteria. Biochem Pharmacol. 2017;133:117–38.

34. Czaplewski L, Bax R, Clokie M, Dawson M, Fairhead H, Fischetti VA, et al. Alternatives to antibiotics-a pipeline portfolio review. Lancet Infect Dis. 2016;16(2):239–51.

35. Kumar P, Kizhakkedathu JN, Straus SK. Antimicrobial Peptides: Diversity, Mechanism of Action and Strategies to Improve the Activity and Biocompatibility In Vivo. Biomolecules. 2018;8(1).

36. Mahlapuu M, Håkansson J, Ringstad L, Björn C. Antimicrobial Peptides: An Emerging Category of Therapeutic Agents. Front Cell Infect Microbiol. 2016;6:194.

37. Bagheri M. Synthesis and thermodynamic characterization of small cyclic antimicrobial arginine and tryptophan-rich peptides with selectivity for Gram-negative bacteria. Methods Mol Biol. 2010;618:87–109.

38. Finger S, Kerth A, Dathe M, Blume A. The efficacy of trivalent cyclic hexapeptides to induce lipid clustering in PG/PE membranes correlates with their antimicrobial activity. Biochim Biophys Acta. 2015;1848(11 Pt A):2998–3006.

39. Gopal R, Na H, Seo CH, Park Y. Antifungal activity of (KW)n or (RW)n peptide against Fusarium solani and Fusarium oxysporum. Int J Mol Sci. 2012;13(11):15042–53.

40. Jakubec M, Rylandsholm FG, Rainsford P, Silk M, Bril’kov M, Kristoffersen T, et al. Goldilocks dilemma: LPS works both as the primary target and a barrier for the antimicrobial action cationic AMPs on E. coli. Doi: 10.20944/preprints202306.2126.v12023.

41. Junkes C, Wessolowski A, Farnaud S, Evans RW, Good L, Bienert M, et al. The interaction of arginine- and tryptophan-rich cyclic hexapeptides with Escherichia coli membranes. J Pept Sci. 2008;14(4):535–43.

42. Rainsford P, Rylandsholm F, Jakubec M, Juskewitz E, Silk M, Engh RA, et al. Label-free measurement of antimicrobial peptide interactions with lipid vesicles and nanodiscs using microscale thermophoresis.. ChemRxiv. 2022.

43. Deschamps E, Schaumann A, Schmitz-Afonso I, Afonso C, Dé E, Loutelier-Bourhis C, et al. Membrane phospholipid composition of Pseudomonas aeruginosa grown in a cystic fibrosis mucus-mimicking medium. Biochim Biophys Acta Biomembr. 2021;1863(1):183482.

44. Wassef MK. Lipids of Klebsiella pneumoniae: the presence of phosphatidyl choline in succinate-grown cells. Lipids. 1976;11(5):364–9.

45. Tao Y, Acket S, Beaumont E, Galez H, Duma L, Rossez Y. Colistin Treatment Affects Lipid Composition of. Antibiotics (Basel). 2021;10(5).

46. Sohlenkamp C, Geiger O. Bacterial membrane lipids: diversity in structures and pathways. FEMS Microbiol Rev. 2016;40(1):133–59.

47. Heung LJ, Luberto C, Del Poeta M. Role of sphingolipids in microbial pathogenesis. Infect Immun. 2006;74(1):28–39.

48. Montefusco DJ, Matmati N, Hannun YA. The yeast sphingolipid signaling landscape. Chem Phys Lipids. 2014;177:26–40.

49. Alvarez HM, Steinbuchel A. Triacylglycerols in prokaryotic microorganisms. Appl Microbiol Biot. 2002;60(4):367–76.

50. Han ML, Zhu Y, Creek DJ, Lin YW, Anderson D, Shen HH, et al. Alterations of Metabolic and Lipid Profiles in Polymyxin-Resistant Pseudomonas aeruginosa. Antimicrob Agents Chemother. 2018;62(6).

51. Schenk ER, Nau F, Thompson CJ, Tse-Dinh YC, Fernandez-Lima F. Changes in lipid distribution in E. coli strains in response to norfloxacin. J Mass Spectrom. 2015;50(1):88–94.

52. Shireen T, Singh M, Dhawan B, Mukhopadhyay K. Characterization of cell membrane parameters of clinical isolates of Staphylococcus aureus with varied susceptibility to alpha-melanocyte stimulating hormone. Peptides. 2012;37(2):334–9.

53. Bisignano C, Ginestra G, Smeriglio A, La Camera E, Crisafi G, Franchina FA, et al. Study of the Lipid Profile of ATCC and Clinical Strains of. Molecules. 2019;24(7).

54. Staubitz P, Neumann H, Schneider T, Wiedemann I, Peschel A. MprF-mediated biosynthesis of lysylphosphatidylglycerol, an important determinant in staphylococcal defensin resistance. FEMS Microbiol Lett. 2004;231(1):67–71.

55. Figueira TN, Freire JM, Cunha-Santos C, Heras M, Gonçalves J, Moscona A, et al. Quantitative analysis of molecular partition towards lipid membranes using surface plasmon resonance. Sci Rep. 2017;7:45647.

56. Hasselmann CaESCM. Determination of minimum inhibitory concentrations (MICs) of antibacterial agents by broth dilution. Clinical Microbiology and Infection. 2003;9(8).

57. Bligh EG, Dyer WJ. A Rapid Method of Total Lipid Extraction and Purification. Can J Biochem Phys. 1959;37(8):911–7.

58. Furse S, Jakubec M, Rise F, Williams HE, Rees CED, Halskau O. Evidence that Listeria innocua modulates its membrane’s stored curvature elastic stress, but not fluidity, through the cell cycle. Sci Rep-Uk. 2017;7.

