## Supplementary_MSmethod_OMVs as bacterial membrane models for "Outer membrane vesicles as realistic models of bacterial membranes in interaction studies by Surface Plasmon Resonance"

```
Thermo Scientific SII for Xcalibur Method

---- Overview ----
Name: New Instrument Method
Comment:
Run time: 32.500 [min]
Instrument: Vanquish_H on thermo-jr7hgt84
Description:
---- Script ----
initial      Instrument Setup
             ColumnComp.PrehtLeft.ReadyTempDelta: 1.00 [°C]
             ColumnComp.PrehtLeft.TempCtrl: On
             ColumnComp.PrehtLeft.Temperature.Nominal: 60.00 [°C]
             ColumnComp.PrehtLeft.EquilibrationTime: 1.0 [min]
             ColumnComp.CC.Mode: StillAir
             ColumnComp.CC.ReadyTempDelta: 0.50 [°C]
             ColumnComp.CC.TempCtrl: On
             ColumnComp.CC.Temperature.Nominal: 60.00 [°C]
             ColumnComp.CC.EquilibrationTime: 1.0 [min]
             ColumnComp.Column_B.ActiveColumn: No
             ColumnComp.Column_B.SystemPressure: "Pump"
             ColumnComp.Column_D.ActiveColumn: Yes
             SamplerModule.Sampler.PunctureOffset: 0 [µm]
             SamplerModule.Sampler.WashSpeed: 20.0 [µl/s]
             SamplerModule.Sampler.InjectWashMode: Both
             SamplerModule.Sampler.WashTime: 5.0 [s]
             SamplerModule.Sampler.DispenseSpeed: 5.000 [µl/s]
             SamplerModule.Sampler.DrawSpeed: 5.000 [µl/s]
             SamplerModule.Sampler.Pump: "Pump"
             SamplerModule.TempCtrl: On
             SamplerModule.Temperature.Nominal: 25.0 [°C]
             PumpModule.Pump.%B_Selector: %B1
             PumpModule.Pump.%A_Selector: %A3
             PumpModule.Pump.%A1_Equate: "Water 1 mM NH4FA 0.01 FA"
             PumpModule.Pump.%A2_Equate: "Water 0.1% FA"
             PumpModule.Pump.%A3_Equate: "50% ACN 1 mM NH4FA 0.01 FA"
             PumpModule.Pump.%B1_Equate: "50/50 IPA/MeCN 1 mM NH4FA 0.01 FA"
             PumpModule.Pump.%B2_Equate: "MeCN 0.1% FA"
             PumpModule.Pump.%B3_Equate: "95% ACN 1 mM NH4FA 0.01 FA"
             PumpModule.Pump.Pressure.LowerLimit: 0 [bar]
             PumpModule.Pump.Pressure.UpperLimit: 1517 [bar]
             PumpModule.Pump.MaximumFlowRampUp: 6.00 [ml/min²]
             PumpModule.Pump.MaximumFlowRampDown: 6.00 [ml/min²]
             ColumnComp.LowerValve.CurrentPosition: 6_1
             ColumnComp.UpperValve.CurrentPosition: 6_1
-2.500 [min] Equilibration
             PumpModule.Pump.Flow.Nominal: 0.600 [ml/min]
             PumpModule.Pump.%B.Value: 30.0 [%]
             PumpModule.Pump.Curve: 5
0.000 [min]
```

```
Thermo Scientific SII for Xcalibur Method

            PumpModule.Pump.Flow.Nominal: 0.600 [ml/min]
            PumpModule.Pump.%B.Value: 30.0 [%]
            PumpModule.Pump.Curve: 5
0.000 [min] Inject Preparation
            Wait ColumnComp.Ready And SamplerModule.Sampler.Ready And PumpModule.Pump.Ready
0.000 [min] Inject
            SamplerModule.Sampler.Inject
0.000 [min] Start Run
            ColumnComp.CC_Temp.AcqOn
            ColumnComp.PrehtLeft_Temp.AcqOn
            PumpModule.Pump.Pump_Pressure.AcqOn
0.000 [min] Run
            PumpModule.Pump.Flow.Nominal: 0.600 [ml/min]
            PumpModule.Pump.%B.Value: 30.0 [%]
            PumpModule.Pump.Curve: 5
20.000 [min]
            PumpModule.Pump.Flow.Nominal: 0.600 [ml/min]
            PumpModule.Pump.%B.Value: 75.0 [%]
            PumpModule.Pump.Curve: 5
25.000 [min]
            PumpModule.Pump.Flow.Nominal: 0.600 [ml/min]
            PumpModule.Pump.%B.Value: 95.0 [%]
            PumpModule.Pump.Curve: 5
30.000 [min] Stop Run
            ColumnComp.CC_Temp.AcqOff
            ColumnComp.PrehtLeft_Temp.AcqOff
            PumpModule.Pump.Pump_Pressure.AcqOff
```

### Method Summary

#### Method Settings

Application Mode: **Small Molecule**
Method Duration (min): **30**

#### Global Parameters

##### Ion Source

Use Ion Source Settings from Tune: **True**
FAIMS Mode: **Not Installed**

##### MS Global Settings

Infusion Mode: **Liquid Chromatography**
Expected LC Peak Width (s): **3**
Advanced Peak Determination: **False**
Mild Trapping: **False**
Default Charge State: **1**
Enable Xcalibur AcquireX method modifications: **True**
Internal Mass Calibration: **EASY-IC™**

##### Divert Valve A

| Time (min) | Position |
| --- | --- |
| 0 | 1-2 |

#### Experiment#1 [AcquireX lipid characterization HCD-CID-MS3]

Start Time (min): **0**
End Time (min): **30**
Cycle Time (sec): **1.5**

##### Master Scan:

### MS OT

Detector Type: **Orbitrap**
Orbitrap Resolution: **120000**
Use Quadrupole Isolation: **True**
Scan Range (m/z): **250-1500**
RF Lens (%): **40**
AGC Target: **Standard**
Maximum Injection Time Mode: **Custom**
Maximum Injection Time (ms): **50**
Microscans: **1**
Data Type: **Profile**
Polarity: **Positive**
Source Fragmentation: **Disabled**
Use EASY-IC™: **True**
Scan Description:

#### Filters:

##### Intensity

Filter Type: **Intensity Threshold**
Intensity Threshold: **1.0e5**

##### Dynamic Exclusion

Exclude after n times: **1**
Exclusion duration (s): **2**
Mass Tolerance: **ppm**
Low: **10**
High: **10**
Exclude Isotopes: **True**

##### Targeted Mass Exclusion

##### Mass List

Mass List Type: **m/z**
Time Mode: **Start/End Time**
Include Intensity Threshold: **True**
Add Mass List Targets Determined by Xcalibur AcquireX: **True**

| Compound | m/z | t start (min) | t stop (min) | Intensity Threshold |
|---|---|---|---|---|
| | 524.265 | 0 | 30 | 1E+20 |

Exclusion mass width: **ppm**
Low: **10**

High: **10**

#### Data Dependent

Data Dependent Mode: **Cycle Time**
Time between Master Scans (sec): **1.5**

#### Scan Event Type 1:

#### Targeted Mass

#### Mass List

Mass List Type: **m/z**
Time Mode: **Start/End Time**
Include Intensity Threshold: **True**
Add Mass List Targets Determined by Xcalibur AcquireX: **True**

| Compound | m/z | t start (min) | t stop (min) | Intensity Threshold |
|---|---|---|---|---|
| | 524.265 | 0 | 30 | 0 |

Mass Tolerance: **ppm**
Low: **10**
High: **10**
Set Collision Energy per Compound: **False**
Perform dependent scan on most intense ion if no targets are found: **True**
Use Group IDs: **False**

#### Scan:

#### ddMS$^2$ OT HCD

Isolation Mode: **Quadrupole**
Isolation Window (m/z): **1.5**
Isolation Offset: **Off**
Activation Type: **HCD**
Collision Energy Mode: **Stepped**
HCD Collision Energy Type: **Normalized**
HCD Collision Energies (%): **25,30,35**
Detector Type: **Orbitrap**
Orbitrap Resolution: **15000**
Scan Range Mode: **Define First Mass**
First Mass (m/z): **140**
AGC Target: **Standard**
Maximum Injection Time Mode: **Custom**

Maximum Injection Time (ms): **50**
Microscans: **1**
Data Type: **Profile**
Use EASY-IC™: **True**
Scan Description:

#### Data Dependent

Data Dependent Mode: **Scans Per Outcome**

#### Scan Event Type 1:

#### Targeted Mass Trigger

#### Mass List

Mass List Type: **m/z**

| Compound | m/z |
|---|---|
| | 184.0733 |

Mass Tolerance: **ppm**
Low: **10**
High: **10**
Use Group IDs: **False**
Trigger Only with Detection of at Least N Ions from the List: **False**
Only Ion(s) Within Top N Most Intense: **True**
n :: **3**
Only Ion(s) Above the Threshold (Relative Intensity, %): **False**
Trigger Type: **Continue Trigger**

#### Intensity

Filter Type: **Intensity Threshold**
Intensity Threshold: **5.0e4**

#### Scan:

#### ddMS$^2$ OT CID

MS$^n$ Level: **2**
Scan Priority: **1**
Isolation Mode: **Quadrupole**
Isolation Window (m/z): **2**
Isolation Offset: **Off**

Activation Type: **CID**
Collision Energy Mode: **Fixed**
CID Collision Energy (%): **32**
CID Activation Time (ms): **10**
Activation Q: **0.25**
Multistage Activation: **False**
Detector Type: **Orbitrap**
Orbitrap Resolution: **15000**
Scan Range Mode: **Auto**
AGC Target: **Standard**
Maximum Injection Time Mode: **Custom**
Maximum Injection Time (ms): **50**
Microscans: **1**
Data Type: **Profile**
Use EASY-IC™: **False**
Scan Description:
Number of Dependent Scans: **1**

#### Scan Event Type 2:

#### Targeted Loss Trigger

#### Mass List

Mass List Type: **m/z**

| Compound | m/z |
|---|---|
| | 217.2042 |
| | 215.1885 |
| | 245.2355 |
| | 243.2198 |
| | 273.2668 |
| | 271.2511 |
| | 269.2355 |
| | 287.2824 |
| | 285.2668 |
| | 301.2981 |
| | 299.2824 |
| | 297.2668 |

| | |
|---|---|
| | 295.2511 |
| | 315.3137 |
| | 329.3294 |
| | 327.3137 |
| | 325.2981 |
| | 323.2824 |
| | 321.2668 |
| | 319.2511 |
| | 343.345 |
| | 357.3607 |
| | 355.345 |
| | 353.3294 |
| | 351.3137 |
| | 349.2981 |
| | 347.2824 |
| | 345.2668 |
| | 371.3763 |
| | 385.392 |
| | 383.3763 |
| | 413.4233 |

Mass Tolerance: **ppm**
Low: **10**
High: **10**
Trigger Only with Detection of at Least N Ions from the List: **False**
Only Ion(s) Within Top N Most Intense: **True**
n :: **3**
Only Ion(s) Above the Threshold (Relative Intensity, %): **False**
Trigger Only with Detection of Correct Charge State of the Product Ion: **False**
Ignore Charge State Requirement for Unassigned Ions: **False**
Trigger Type: **Continue Trigger**

##### Precursor Ion Exclusion

Exclusion mass width: **m/z**
Low: **0.5**
High: **4**

#### Intensity

Filter Type: **Intensity Threshold**
Intensity Threshold: **5.0e4**

#### Scan:

#### ddMS$^3$ OT CID

MS$^n$ Level: **3**
Scan Priority: **1**
MS Isolation Window (m/z): **1.5**
MS2 Isolation Window (m/z): **2**
Isolation Offset: **Off**
Activation Type: **CID**
Collision Energy Mode: **Fixed**
CID Collision Energy (%): **35**
CID Activation Time (ms): **10**
Activation Q: **0.25**
Multistage Activation: **False**
Detector Type: **Orbitrap**
Orbitrap Resolution: **15000**
Scan Range Mode: **Auto**
AGC Target: **Standard**
Maximum Injection Time Mode: **Custom**
Maximum Injection Time (ms): **65**
Microscans: **1**
Data Type: **Profile**
Use EASY-IC™: **False**
Scan Description:
Number of Dependent Scans: **3**
