## Supplementary_tablesandfigure_OMVs as bacterial membrane models for "Outer membrane vesicles as realistic models of bacterial membranes in interaction studies by Surface Plasmon Resonance"

### SUPPLEMENTARY TABLES AND FIGURES

#### Table S1

Table S1. Characterization of size and zeta potential of the isolated OMVs and liposomes prepared from *E. coli* Δ*tolA* lipid isolate.

| Sample | Size, nm | PDI | Zeta potential, mV | |
|---|---|---|---|---|
| | | | OMVs | Cell |
| Liposomes | 139.9 ± 1.6 | 0.299 ± 0.01 | − 17.80 ± 0.5 | |
| OMVs: | | | | |
| *E. coli* Δ*tolA* | 82.7 ± 1.3 | 0.246 ± 0.01 | − 9.4 ± 0.8 | − 6.07 ± 1.4 |
| *K. pneumoniae* | 194.8 ± 1.0 | 0.333 ± 0.01 | − 9.9 ± 0.3 | − 5.39 ± 0.8 |
| *A. baumannii* | 175.0 ± 1.3 | 0.240 ± 0.00 | − 14.4 ± 0.6 | − 9.20 ± 0.5 |
| *P. aeruginosa* | 143.3 ± 1.1 | 0.103 ± 0.01 | − 3.90± 0.6 | − 4.71 ± 1.6 |

#### Table S2

Table S2. Fatty acid composition within PE class of lipids analyzed by mass spectrometry

| Strains | PE fatty acid composition | Lipid isolate | | PE fatty acid composition | OMVs | |
|---|---|---|---|---|---|---|
| | | Relative abundance | st. dev. | | Relative abundance | st. dev. |
| *E. coli ΔtolA* | PE 16:0_16:1 | 0.19 | 0.04 | PE 16:0_16:1 | 0.27 | 0.09 |
| | PE 16:1_18:1 | 0.16 | 0.00 | PE 16:1_18:1 | 0.12 | 0.06 |
| | PE 16:0_18:1 | 0.12 | 0.01 | PE 16:0_18:1 | 0.11 | 0.01 |
| | PE 18:1_18:1 | 0.11 | 0.04 | PE 16:0_17:1 | 0.08 | 0.02 |
| | PE 16:0_17:1 | 0.09 | 0.00 | PE 18:1_18:1 | 0.06 | 0.01 |
| | PE 18:4_16:0 | 0.03 | 0.00 | PE 18:4_16:0 | 0.05 | 0.00 |
| | PE 17:1_18:1 | 0.03 | 0.03 | PE 16:1_16:1 | 0.04 | 0.00 |
| | PE 16:1_16:1 | 0.02 | 0.00 | PE 14:0_16:0 | 0.03 | 0.00 |
| | PE 20:4_16:1 | 0.02 | 0.00 | PE 17:1_18:1 | 0.02 | 0.00 |
| | PE 16:1_17:1 | 0.02 | 0.00 | PE 20:4_16:1 | 0.02 | 0.00 |
| | PE 14:0_16:0 | 0.02 | 0.00 | PE 18:3_16:1 | 0.02 | 0.01 |
| *K. pneumoniae* | PE 16:0_17:1 | 0.29 | 0.04 | PE 16:0_16:1 | 0.12 | 0.01 |
| | PE 16:0_19:1 | 0.16 | 0.00 | PE 14:0_16:0 | 0.12 | 0.03 |
| | PE 14:0_16:0 | 0.15 | 0.01 | PE 16:0_17:1 | 0.12 | 0.00 |
| | PE 19:1_17:1 | 0.10 | 0.00 | PE 16:0_18:1 | 0.12 | 0.01 |
| | PE 19:1_19:1 | 0.05 | 0.01 | PE 16:0_19:1 | 0.08 | 0.00 |
| | PE 14:0_17:1 | 0.05 | 0.00 | PE 16:0_16:0 | 0.04 | 0.00 |
| | PE 17:1_17:1 | 0.03 | 0.00 | PE 16:1_18:1 | 0.04 | 0.01 |
| | PE 16:0_16:0 | 0.03 | 0.01 | PE 14:0_17:1 | 0.03 | 0.00 |
| | PE 14:0_14:0 | 0.01 | 0.01 | PE 16:0_18:1 | 0.02 | 0.01 |
| *A. baumannii* | PE 16:0_18:1 | 0.28 | 0.03 | PE 16:0_18:1 | 0.35 | 0.03 |
| | PE 18:1_18:1 | 0.16 | 0.00 | PE 16:1_18:1 | 0.25 | 0.00 |
| | PE 16:1_18:1 | 0.10 | 0.01 | PE 18:0_18:1 | 0.04 | 0.00 |
| | PE 14:0_18:1 | 0.09 | 0.00 | PE 18:1_18:0 | 0.04 | 0.00 |
| | PE 15:0_18:1 | 0.06 | 0.00 | PE 18:1_18:1 | 0.04 | 0.01 |
| | PE 16:0_19:1 | 0.06 | 0.00 | PE 16:0_16:1 | 0.04 | 0.00 |
| | PE 16:0_16:0 | 0.05 | 0.00 | PE 16:0_16:0 | 0.02 | 0.00 |
| | PE 17:1_18:1 | 0.02 | 0.00 | PE 16:1_16:1 | 0.02 | 0.00 |
| | PE 18:4_19:0 | 0.02 | 0.00 | PE 16:0_19:1 | 0.01 | 0.00 |
| *P. aeruginosa* | PE 16:0_18:1 | 0.36 | 0.07 | PE 16:0_18:1 | 0.44 | 0.01 |
| | PE 16:0_19:1 | 0.13 | 0.00 | PE 16:1_18:1 | 0.11 | 0.03 |
| | PE 16:1_18:1 | 0.09 | 0.00 | PE 16:0_16:1 | 0.06 | 0.00 |
| | PE 17:1_18:1 | 0.04 | 0.00 | PE 16:0_16:0 | 0.06 | 0.00 |
| | PE 15:0_18:1 | 0.04 | 0.00 | PE 16:0_18:0 | 0.04 | 0.00 |
| | PE 16:1_18:1 | 0.04 | 0.01 | PE 16:0_16:1 | 0.02 | 0.00 |
| | PE 16:0_16:0 | 0.04 | 0.00 | PE 18:1_18:1 | 0.02 | 0.00 |
| | PE 14:0_18:1 | 0.03 | 0.01 | PE 16:1_18:1 | 0.02 | 0.01 |
| | PE 18:1_18:1 | 0.03 | 0.01 | PE 16:0_19:1 | 0.01 | 0.00 |

#### Table S3

Table S3. Dissociation constant $K_D$ determined by SPR

| AMP | $K_D$, μM | | | | |
|---|---|---|---|---|---|
| | Liposomes | *E. coli ΔtolA* | *K. pneumoniae* | *A. baumannii* | *P. aeruginosa* |
| c-LWwNKr | 966.5 ± 170.1 | 406.4 ± 38.1 | 1075.2 ± 49.0 | 857.8 ± 61.4 | 1036.5 ± 269.7 |
| c-WKWKWK | 244.4 ± 39.5 | 380.9 ± 22.6 | 693.7 ± 37.3 | 389.0 ± 32.5 | 749.3 ± 117.3 |
| c-WRWRWR | 74.5 ± 3.3 | 73.0 ± 7.3 | 81.1 ± 2.2 | 113.6 ± 5.2 | 118.5 ± 3.5 |
| c-WWWKKK | 28.3 ± 4.2 | 56.5 ± 1.0 | 21.0 ± 0.7 | 100.2 ± 13.1 | 100.0 ± 4.1 |
| c-WWWRRR | 47.8 ± 5.2 | 27.0 ± 1.6 | 35.8 ± 1.3 | 48.7 ± 2.7 | 59.0 ± 2.6 |

#### Table S4

Table S4. $k_{off}$ rate constant determined by SPR

| AMP | $k_{off}$, $s^{-1}$ | | | | |
|---|---|---|---|---|---|
| | Liposomes | *E. coli ΔtolA* | *K. pneumoniae* | *A. baumannii* | *P. aeruginosa* |
| c-LWwNKr | 1.67 ± 0.45 | 1.42 ± 0.27 | 1.75 ± 0.55 | 1.72 ± 0.45 | 1.78 ± 0.54 |
| c-WKWKWK | 0.95 ± 0.20 | 1.42 ± 0.16 | 1.51 ± 0.54 | 0.82 ± 0.11 | 1.52 ± 0.37 |
| c-WRWRWR | 0.57 ± 0.09 | 0.74 ± 0.10 | 0.59 ± 0.10 | 0.64 ± 0.13 | 0.42 ± 0.07 |
| c-WWWKKK | 0.21 ± 0.01 | 0.62 ± 0.05 | 0.37 ± 0.03 | 0.63 ± 0.12 | 0.62 ± 0.07 |
| c-WWWRRR | 0.51 ± 0.08 | 0.18 ± 0.02 | 0.28 ± 0.01 | 0.40 ± 0.06 | 0.37 ± 0.04 |

#### Table S5

Table S5. Calculated $k_{on}$ rate constant

| AMP | $k_{on}$, $\mu M^{-1} \cdot s^{-1}$ | | | | |
|---|---|---|---|---|---|
| | Liposomes | *E. coli ΔtolA* | *K. pneumoniae* | *A. baumannii* | *P. aeruginosa* |
| c-LWwNKr | 0.002 ± 0.001 | 0.003 ± 0.001 | 0.002 ± 0.001 | 0.002 ± 0.001 | 0.002 ± 0.001 |
| c-WKWKWK | 0.004 ± 0.001 | 0.004 ± 0.001 | 0.002 ± 0.001 | 0.002 ± 0.001 | 0.002 ± 0.001 |
| c-WRWRWR | 0.008 ± 0.001 | 0.010 ± 0.001 | 0.007 ± 0.001 | 0.006 ± 0.001 | 0.004 ± 0.001 |
| c-WWWKKK | 0.007 ± 0.001 | 0.011 ± 0.001 | 0.018 ± 0.001 | 0.006 ± 0.002 | 0.006 ± 0.001 |
| c-WWWRRR | 0.011 ± 0.001 | 0.007 ± 0.001 | 0.008 ± 0.001 | 0.008 ± 0.002 | 0.006 ± 0.001 |

### Table S6

Table S6. Summary of statistical analysis of SPR data with T-test

| AMP | p-value | $K_D$ | | | | $k_{off}$ | | | | $k_{on}$ | | | |
|---|---|---|---|---|---|---|---|---|---|---|---|---|---|
| | | LI | EC | KP | AB | LI | EC | KP | AB | LI | EC | KP | AB |
| c-LWwNKr | EC | 0.00 | – | – | – | 0.22 | – | – | – | 0.01 | – | – | – |
| | KP | 0.17 | 0.00 | – | – | 0.43 | 0.20 | – | – | 0.40 | 0.01 | – | – |
| | AB | 0.18 | 0.00 | 0.00 | – | 0.45 | 0.19 | 0.47 | – | 0.32 | 0.02 | 0.24 | |
| | PA | 0.36 | 0.01 | 0.41 | 0.16 | 0.40 | 0.18 | 0.47 | 0.45 | 0.43 | 0.00 | 0.43 | 0.22 |
| c-WKWKWK | EC | 0.00 | – | – | – | 0.02 | – | – | – | 0.38 | – | – | – |
| | KP | 0.00 | 0.00 | – | – | 0.05 | 0.40 | – | – | 0.01 | 0.02 | – | – |
| | AB | 0.00 | 0.37 | 0.00 | | 0.19 | 0,00 | 0.05 | – | 0.00 | 0.01 | 0.48 | – |
| | PA | 0.00 | 0.00 | 0.24 | 0.00 | 0,03 | 0.26 | 0.43 | 0.01 | 0.00 | 0.01 | 0.46 | 0.46 |
| c-WRWRWR | EC | 0.38 | – | – | – | 0.05 | – | – | – | 0.03 | – | – | – |
| | KP | 0.02 | 0.07 | – | – | 0.41 | 0.07 | – | – | 0.37 | 0.02 | – | – |
| | AB | 0.00 | 0.00 | 0.00 | – | 0.22 | 0.18 | 0.29 | – | 0.07 | 0.00 | 0.01 | – |
| | PA | 0.00 | 0.00 | 0.00 | 0.24 | 0.04 | 0.01 | 0.04 | 0.03 | 0.01 | 0.00 | 0.01 | 0.02 |
| c-WWWKKK | EC | 0.00 | – | – | – | 0.00 | – | – | – | 0.01 | – | – | – |
| | KP | 0.02 | 0.00 | – | – | 0.00 | 0.00 | – | – | 0.00 | 0.00 | – | – |
| | AB | 0.00 | 0.00 | 0.00 | – | 0.00 | 0.41 | 0.01 | – | 0.26 | 0.01 | 0.00 | – |
| | PA | 0.00 | 0.00 | 0.00 | 0.49 | 0.00 | 0.48 | 0.00 | 0.43 | 0.13 | 0.00 | 0.00 | 0.41 |
| c-WRWRWR | EC | 0.00 | – | – | – | 0.00 | – | – | – | 0.03 | – | – | – |
| | KP | 0.01 | 0.00 | – | – | 0.00 | 0.00 | – | – | 0.06 | 0.05 | – | – |
| | AB | 0.40 | 0.00 | 0.00 | – | 0.01 | 0.00 | 0.01 | – | 0.06 | 0.05 | 0.23 | – |
| | PA | 0.01 | 0.00 | 0.00 | 0.00 | 0.02 | 0.00 | 0.01 | 0.03 | 0.02 | 0.33 | 0.03 | 0.00 |

Here: LI – liposomes prepared from *E. coli* Δ*tolA* lipid isolate, EC – *E. coli* Δ*tolA*, KP – *K. pneumoniae*, AB – *A. baumannii*, PA – *P. aeruginosa*. p-value ≥ 0.05 is highlighted in red.

#### Figure S1

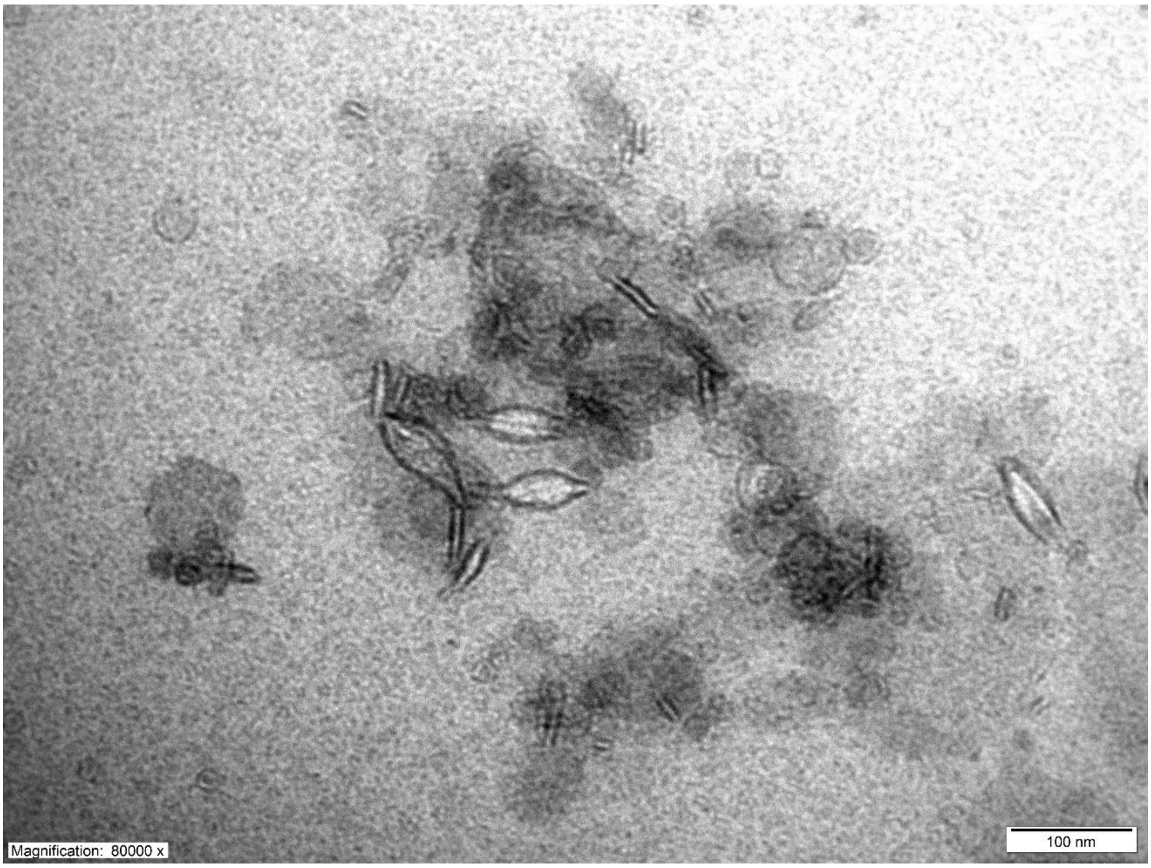


Figure S1. TEM micrograph of OMVs secreted by *E. coli* NR698 strain at 80,000x magnification. 100 nm scalebar is presented at the bottom right corner of the micrograph.
